# Mechanism for Vipp1 spiral formation, ring biogenesis and membrane repair

**DOI:** 10.1101/2023.09.26.559607

**Authors:** Souvik Naskar, Andrea Merino, Javier Espadas, Jayanti Singh, Aurelien Roux, Adai Colom, Harry H Low

## Abstract

ESCRT-III family proteins build dynamic filaments that remodel membrane. Although the transition of planar filaments into 3D membrane budding structures is fundamental for their function, the geometric changes in polymer architecture driving the transition remain obscure. Here we show how bacterial Vipp1 polymerises into dynamic planar sheets and spirals on membrane. The spirals converge to form a central ring like those known to bud membrane. To probe how Vipp1 morphs between polymers, we determine the architecture of multiple helical filaments. As well as describing filament constriction and membrane tubulation, the geometric relationship between helical and planar lattices enables Vipp1 sheets and spirals to be modelled. Moreover, the helical structures show filaments twisting – a process needed for Vipp1 to transition between planar and 3D architectures. Given the structural conservation between Vipp1 and ESCRT-III, our results may represent the broad changes in geometry required for some ESCRT-III filaments to switch between 2D and 3D forms.

## Introduction

ESCRT-III (endosomal sorting complex required for transport-III) family members are ancient membrane remodelling devices with an evolutionary lineage that traces back to LUCA, the last universal common ancestor of cells^1^. Over time the family has radiated across the tree of life acquiring often essential and conserved functions. In eukaryotes and archaea, ESCRT-III systems drive membrane abscission during cell division^2^, promote viral replication and budding^3,4^, and mediate extracellular vesicle biogenesis^5,6^. Other important functions specific to eukaryotes include multivesicular body biogenesis^7^ and membrane repair^8^. In bacteria, where PspA and its paralogue Vipp1 (also IM30) were recently discovered as ESCRT-III homologues^1,9^, these proteins function in membrane stress response and repair. PspA is triggered by agents that threaten the inner membrane including phage, mislocalised secretins and antibiotics^10–14;^ whilst Vipp1 is a universal plastid component in cyanobacteria, algae, and plants where it undertakes essential functions in thylakoid membrane biogenesis and repair^15–24^.

ESCRT-III family members have a conserved core fold consisting of five helices, α1-α5^1,25^. Whilst helices α1 and α2 form a characteristic hairpin motif, in some systems helices α3-α5 switch conformation between open, intermediate, and closed^26^. Some ESCRT-III family members such as Vipp1, Vps2/CHMP2, Vps24/CHMP3, and Snf7/CHMP4 supplement the core fold with an N-terminal motif or amphipathic helix (termed helix α0) that mediates membrane binding^13,27–29^. C-terminal to helix α5 are less conserved elements^1^ that mediate protein interactions in most eukaryotic ESCRT-III systems^25^. In this region, Vipp1 has a ∼40 amino acid C-terminal domain (CTD) that is flexible and may incorporate helix α6^30^. The CTD tunes Vipp1 polymerisation dynamics both *in-vivo* and *in-vitro*^1,30,31^ and might constitute a second lipid binding domain that promotes membrane fusion^31,32^. Using the core fold as a building block, ESCRT-III family members assemble conserved filaments characterised by the hairpin motif of neighbouring subunits stacking side-by-side with helix α5 binding in a domain swop across the hairpin tip^1,7,9,33,34^. This filament is used to build different supramolecular structures including spirals^33,35–44^, helical filaments^9,33,43,45^ and dome-shaped rings^1,34^. In bacteria, although *Synechocystis* Vipp1^46^ and PspA^9^ form planar patches, spiral filaments have not been reported, which currently represents a key differentiating factor from their eukaryotic counterparts.

The assembly of ESCRT-III filaments is fundamental to their membrane remodelling mechanism with the formation of planar spirals on the membrane surface an early key step. Current models describe spirals as loaded springs with elastic stress accumulating due to a preferred radius of curvature. Stress is highest at the spiral perimeter and centre where the filament is under-curved or over-curved, respectively. This stress, which constitutes an energy store, is theoretically sufficient to bend membrane^40,47^. Energy minimisation, through the buckling of planar spiral filaments to 3D polymers such as conical spirals or helices, is directly coupled to the mechanical shaping of bound membrane. An important component of the model depends on the sequential tilting of polymer orientation relative to the membrane^48,49^, with exposure to tilted membrane binding interfaces within the filament a driver of membrane deformation^49,50^. Notably, by simply switching the position of the membrane binding interface to be oriented towards the inside or outside of the tilted filament, the direction of membrane budding can theoretically be reversed with the filament binding and exerting force from the outside or inside of the membrane, respectively^48^. How ESCRT-III subunits are arranged within these planar or twisted filaments when bound to membrane remains unclear.

The structure of Vipp1 when assembled as a dome-shaped ring, combined with the observation that these rings are sufficient to bud membrane, represented an alternative mechanism for how ESCRT-III family members remodel membrane. Vipp1 rings recruited to the surface of both monolayers and precurved lipid bilayers bud the membrane by internalising it within the ring lumen. This movement is mediated by membrane-binding domains (helix α0) lining the inner lumen so that membrane is drawn in via a capillary action-like mechanism. Although Vipp1 and PspA have the capacity to bind and hydrolyse nucleotides^34,55^, this mode of ring-mediated membrane remodelling occurs passively without chemical energy turnover, at least *in-vitro*. The capacity of Vipp1 rings to bud membrane independently of spiral springs hints that a similar process may be an unrecognised contributory factor in eukaryotic ESCRT-III membrane remodelling processes where rings are assembled, albeit transiently, in the centre of spirals^51^.

The biogenesis pathway for Vipp1 rings is unknown. It is also unclear how Vipp1 rings, as well as other polymers such as helical filaments, might relate structurally and functionally to membrane bound planar forms that stabilise and reduce proton permeability in liposomes^46^. These planar forms may be fundamental for how Vipp1 and PspA function in membrane stabilisation and repair across bacteria. Here we use a fusion of light microscopy, fast-atomic force microscopy (F-AFM) and electron microscopy (EM) to show how Vipp1 rings originate from a planar spiral progenitor. By assembling and comparing structural models for different types of Vipp1 polymer including planar sheets, spiral filaments, helices and rings, we suggest a model for Vipp1 ring biogenesis that has implication for how other ESCRT-III systems bud membrane.

## Results

### Vipp1 is a membrane sensor recruited to highly curved and perturbed membranes

Vipp1 from *Nostoc punctiforme* was purified in a low salt (10 mM NaCl) buffer (Figure 1A). This yielded various 3D polymers including helical filaments, helical-like ribbons, and some dome-shaped rings^1^. This Vipp1 sample was used throughout this study unless otherwise specified. To study Vipp1 dynamics on membrane, Vipp1 was labelled with Alexa488 (termed Vipp1_Alexa448_), introduced into a flow chamber incorporating supported lipid bilayers (SLBs; Figure 1B), and imaged by confocal microscopy. This methodology was previously used to characterise how the ESCRT-III protein Snf7 interacted with model membranes^40^. Whilst Snf7 formed evenly distributed patches on the membrane surface, Vipp1_Alexa448_ formed a localised coating at the SLB edge (Figure 1C). Here, the membrane was highly curved with regions of poorly packed or perturbed lipid expected. Shifting to buffer conditions with 500 mM NaCl promoted the partial disassembly of polymers (Figure 1A) and reduced larger soluble fluorescent foci and background fluorescence. However, Vipp1_Alexa448_ still targeted the highly curved SLB edge (Figure 1C-1E). Where neighbouring SLBs fused and coated leading edges were flattened at the membrane merge point, bound Vipp1_Alexa448_ rapidly disassembled from the membrane supporting the notion of a highly dynamic interaction with the membrane edge (Figure 1F). In summary, this data revealed Vipp1_Alexa488_ to have a strong preference for highly curved and perturbed membrane and was consistent with Vipp1 functioning as a membrane curvature sensor.

**Figure 1.**
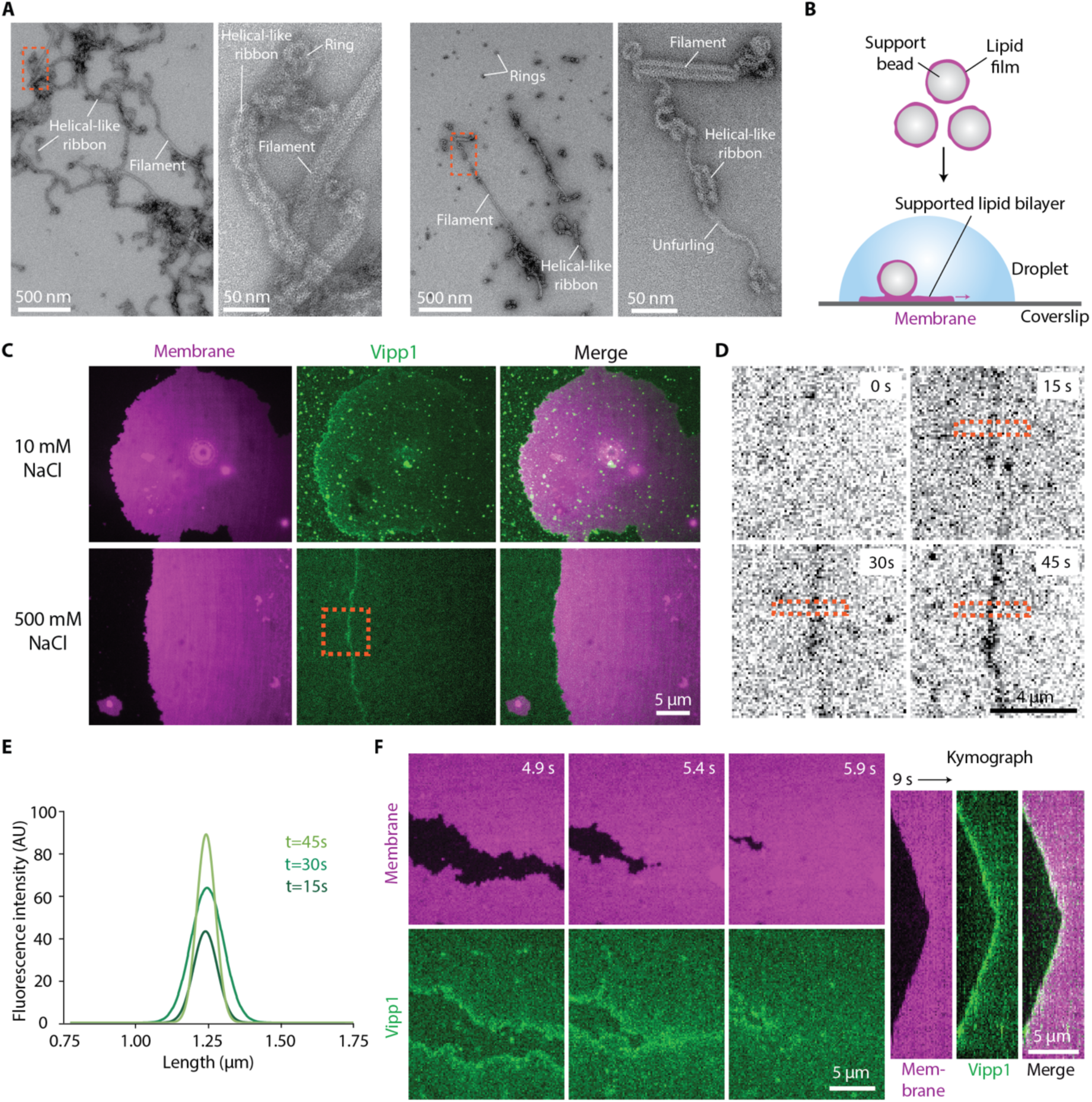
Vipp1 is a membrane sensor recruited to highly curved and perturbed membranes. **A**, Left panel pair with NS-EM image showing Vipp1 purified in low salt (10 mM NaCl) buffer. Neighbouring panel represents zoomed dotted box. Right panel pair with Vipp1 in a 500 mM NaCl buffer. Neighbouring panel represents zoomed dotted box. Note the unfurling of the helical-like ribbon. **B**, Preparation of supported lipid bilayers (SLBs). **C**, Fluorescent microscopy showing Vipp1_Alexa488_ recruitment to the highly curved membrane edge. Vipp1_Alexa488_ recruitment is unaffected by ionic strength. **D**, Timecourse showcasing dynamic Vipp1_Alexa488_ recruitment to the membrane edge. Relates to dotted box in **C**. **E**, Fitted curves of the fluorescence plot profile show increasing Vipp1_Alexa488_ recruitment to the membrane edge. Measurements from dotted box in **D**. **F**, Timecourse with accompanying kymograph (right panel) showing fusion of neighbouring SLBs with Vipp1_Alexa488_ lost from the membrane merge point.

### Vipp1 assembles dynamic networks of spirals, rings and sheets on membrane

To further resolve the dynamics and architecture of Vipp1 on the membrane edge, we utilised F-AFM which provided high temporal and spatial resolutions. Consistent with the fluorescent microscopy data (Figure 1C-1F), Vipp1 accumulated at the edge of SLB patches, again indicating a sensing capability for highly curved or perturbed membrane (Figure 2A). However, we did not observe 3D polymers such as helical filaments (Figure 1A) binding as expected. Instead, Vipp1 grew as dynamic planar filaments that curled anticlockwise to form spirals or sometimes rings (Figures 2B, 2C, Supplementary Figure 1A and Videos S1-S2). These filaments grew from small oligomers (or monomers) that originated from disassembled 3D polymers given the sample was originally gel filtrated. Filaments grew into unpopulated membrane regions so that ultimately networks of planar sheets, spiral filaments and rings covered the entire lipid surface within minutes. Sometimes filaments split indicating a protofilament substructure or nascent filaments grew alongside established filaments to form planar sheets (Supplementary Figures 1B and 1C). Conversely, sheets separated branching off filaments that curled into spirals (Figure 2D and Supplementary Figure 1C). Membrane height offset for both sheet and spiral filaments was ∼5.5 nm suggesting a close structural relationship (Figure 2E). Filaments grew at a mean rate of 24 ± (a standard deviation of) 19.6 nm/s, N=124 independent measurements (Figure 2F) and with generally uniform width with mean 13.4 ± 0.9 nm, N=13 (Figure 2G). Spiral mean diameter was 82.7 nm ± 37.8, N=276 (Figure 2H) with a mean area of 6485 ± 6303 nm^2^, N=276 (Figure 2I). These spirals were reminiscent of Snf7 spirals^40^ although when packed they did not deform into polygons like Snf7, indicating an increased stiffness for the Vipp1 filament. They also did not fragment towards the spiral perimeter as was characteristic of Snf7. Spirals curled in either Archimedes or exponential forms towards their centre until filaments reached a curvature limit beyond which further curling was impeded resulting in filament merging and formation of closed rings (Figures 2D and 3A). This spiral maturation and ring biogenesis correlated with increasing height offset between spiral inner turns and the membrane (Figures 3B, Supplementary Figure 1A and Video S3). In mature spirals, rings protruded 1.0 ± 0.2 nm (N=16) above surrounding spiral filaments (Figure 3C) and eventually detached from the parent spiral (Figure 3A and 3D, Supplementary Figure 1A). These rings, that originated from spirals under low <u>s</u>alt buffer conditions (10 mM NaCl) and with mean diameter 37.0 ± 3.9 nm (N=39) were termed Vipp1 rings_LS_ (Figure 3E and 3F). Periodically, rings formed spontaneously on the curved edge of SLBs or on isolated lipid micro-patches in the absence of a parent spiral (Supplementary Figure 1A, 1D and 1E). Here, the nascent Vipp1 filament grew spatially constrained by the membrane support or surrounding planar filaments. In these instances, mean ring height was 6.0 ± 0.6 nm, N=22, which was similar to Vipp1 rings_LS_ (Supplementary Figure 1E). However, mean diameter was generally wider 49.1 ± 7.8 nm (N=39) than Vipp1 rings_LS_ (Figure 3E and 3F) showing how maximum bending curvature was not generally achieved without the corralling effect of a spiral.

**Figure 2.**
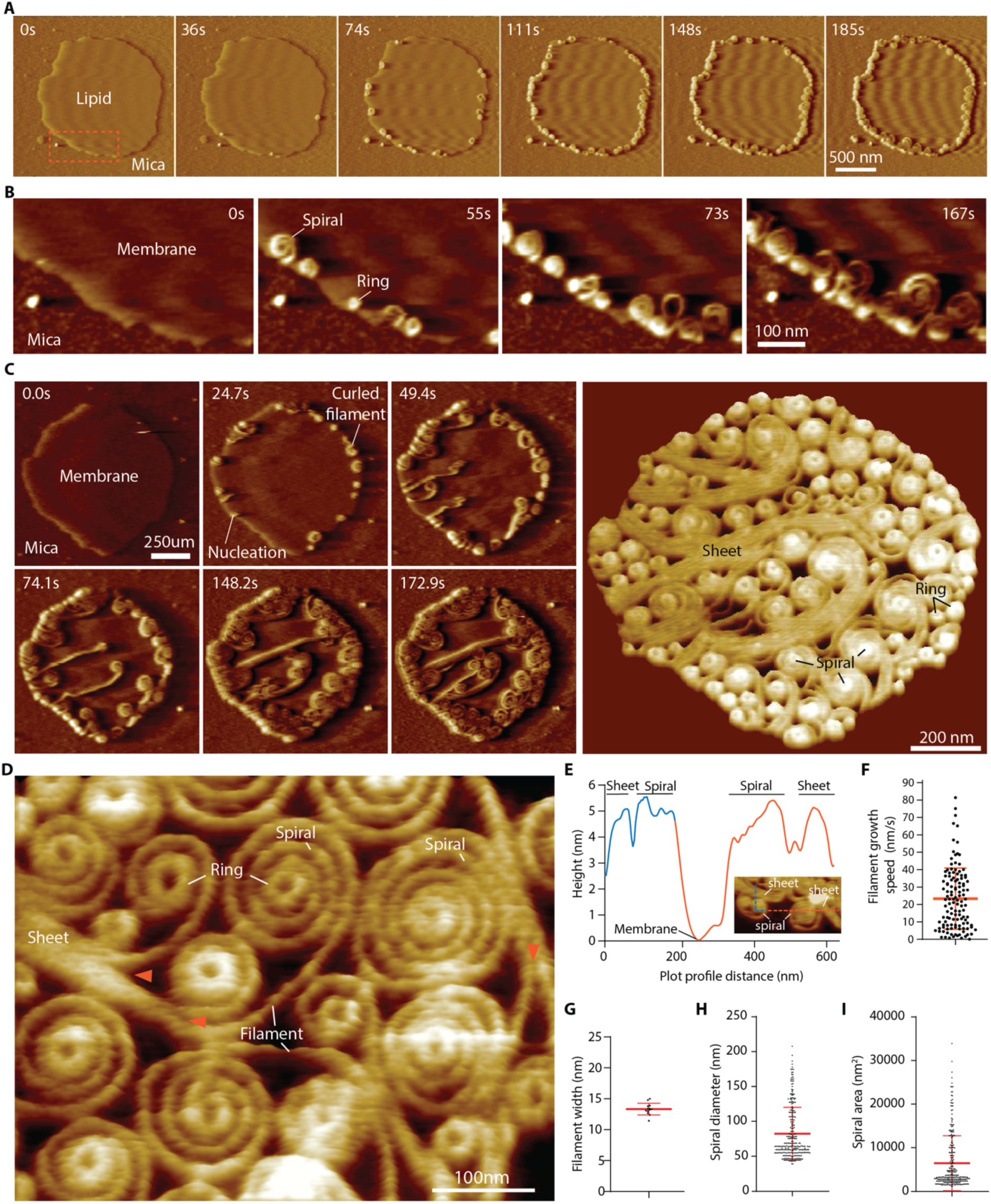
Vipp1 assembles dynamic networks of spirals, rings and sheets on membrane. **A**, F-AFM phase timecourse showing Vipp1 recruitment to the highly curved edge of membrane patches. **B**, Zoom of dotted box in **A** revealing spiral and ring formation localised to the membrane edge. **C**, Phase timecourse showcasing a dense network of sheets, spirals and rings that ultimately cover the entire membrane plane. **D**, Phase image showing Vipp1 sheet, spiral and ring detail. Red arrows indicate sheet branching into filaments ∼13 nm wide. **E**, Vipp1 sheet and spiral filament height offset from the membrane. **F-I**, Quantification of Vipp1 filament and spiral characteristics.

**Figure 3.**
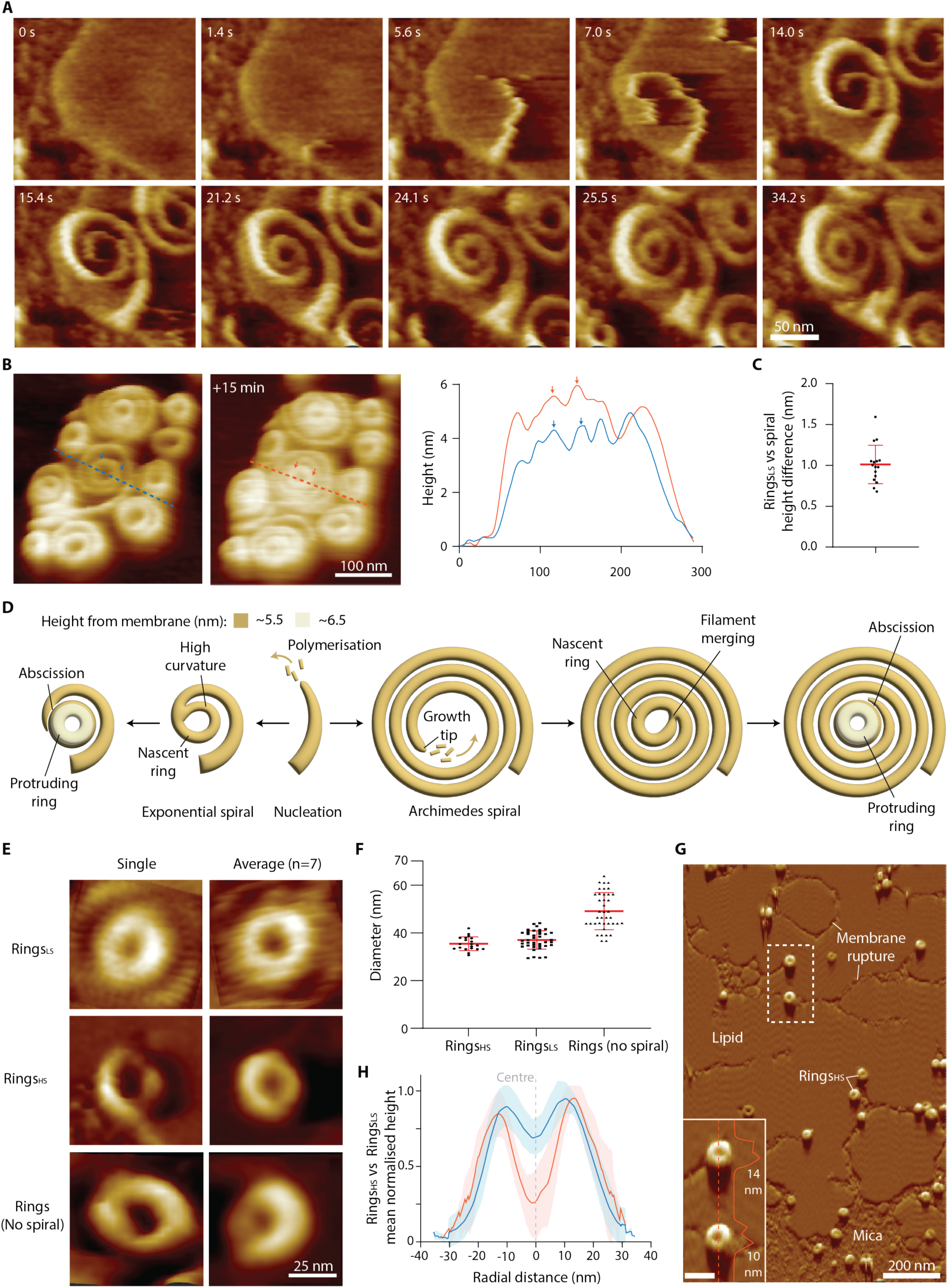
Vipp1 spirals form protruding central rings that abscise. **A**, F-AFM phase timecourse showing ring biogenesis from an exponential-shaped spiral. Relates to Supplementary Figure 1A. **B**, F-AFM timecourse showing how spiral and ring maturation correlates with increased filament offset from the membrane. Blue and red dotted lines indicate plotted height profile. **C**, Quantification of height difference between Vipp1 rings_LS_ and surrounding spiral filaments. **D**, Schematic showing spiral and ring biogenesis pathways. **E**, F-AFM phase images showcasing the different types of Vipp1 ring observed. **F**, Quantification of Vipp1 ring diameters as shown in **E**. **G**, F-AFM phase image where Vipp1 rings_HS_ scan and stably bind highly curved or ruptured membrane. Dotted box indicates zoom panel with 40 nm scale bar. **H**, Quantification of Vipp1 rings_HS_ and rings_LS_ height profiles show similar shape and lateral dimensions.

### Vipp1 rings scan and bind damaged membrane

Previously, dome-shaped rings were purified and structurally characterised revealing a mix of core symmetries ranging from C11 to C17 with diameters from 24 nm to 34 nm, respectively^1^. Larger well-ordered C20 symmetry rings with 41 nm diameter were also prevalent in the sample with the largest abundant ring class ∼43 nm diameter (Supplementary Figure 2A). To obtain these pre-assembled rings, the sample was purified in a <u>h</u>igh <u>s</u>alt (50 mM NaCl) buffer^1^. Here, we termed these rings Vipp1 rings_HS_ to distinguish them from Vipp1 rings_LS_. Given the potential similarity between Vipp1 rings_HS_ and Vipp1 rings_LS_, we aimed to characterise Vipp1 rings_HS_ by F-AFM. Vipp1 rings_HS_ exposed to SLBs bound the membrane surface and maintained their pre-assembled ring structure (Figure 3G and Supplementary Figure 2B). Intriguingly, Vipp1 rings_HS_ targeted the highly curved edge of membrane patches or rupture lines in the lipid. Vipp1 rings_HS_ therefore have the remarkable capability of scanning membrane surfaces for damaged regions before targeting them for binding. By F-AFM, Vipp1 rings_HS_ had a mean diameter of 35.5 ± 2.9 nm, N=20 (Figure 3E and 3F), which correlated well with the mean diameter of Vipp1 rings_LS_ and the larger ring diameters observed by EM (Supplementary Figure 2A)^1^. Vipp1 rings_HS_ had a mean height of 9.6 ± 2.2 nm, N=46 (Supplementary Figure 2C), which was ∼3 nm higher than Vipp1 rings_LS_ but markedly lower than cryo-EM determined Vipp1 ring_HS_ heights between ∼15-21 nm for C11-C17 symmetries^1^. For the latter, the height difference was likely due to disassembly of the lower rungs of the dome-shaped rings into the membrane once bound, although compression from the F-AFM tip cannot be discounted. Overall, Vipp1 rings_HS_ function as membrane sensing scaffolds and are broadly similar in shape and form to Vipp1 rings_LS_ when measured by F-AFM (Figure 3H).

Next we examined the effect of low salt (10 mM NaCl) on Vipp1 rings_HS_. Intriguingly, Vipp1 rings_HS_ were no longer observed binding the lipid. Instead, they disassembled and nucleated spirals, sheets and rings that grew from the membrane edge (Supplementary Figure 2D and 2E). Negative stain EM verified the polymeric state of Vipp1 rings_HS_ in the low salt buffer (Supplementary Figure 2F) although rings had increased tendency to clump or unfurl into quasi-helical ribbons showcasing the effect of low salt on destabilising the rings and promoting a transition towards helical-like filaments.

### Vipp1 spiral filaments and planar sheets have closely related lattices

To analyse the relationship between Vipp1 sheet and spiral filament ultrastructure under low salt conditions (10 mM NaCl), lower velocity AFM images were obtained which revealed remarkable detail. Sheets were highly ordered forming parallel stripes or ridges spaced 54 Å apart (Figure 4A). Orthogonal to the ridge lines, parallel seams could be discerned in the sheets spaced 122 Å apart, which was consistent with the width of the filaments in spiral turns (Figure 2G). This supports a model where sheets form planar crystalline super structures comprised of merged filaments ∼12-13 nm across and are predisposed to branch and spiral at the seam points. Analysis of spiral filaments also revealed parallel ridges spaced 54 Å apart (Figure 4B). Neighbouring spiral turns sometimes merged indicating filaments may transition into localised sheet patches under different conditions. To further resolve Vipp1 planar ultrastructure, Vipp1 was exposed to lipid monolayers. In contrast to the SLB system where Vipp1 3D polymers did not bind directly to the membrane (apart from Vipp1 rings_HS_), 3D polymers such as rod-like filaments (∼35 nm diameter) were sometimes observed bound to the lipid monolayer surface (Figure 4C). These presumably originated as ordered helical filaments in the initial sample (Figure 1A). They often had tips that appeared to be flattening and merging with the surrounding milieu. Remarkably, close inspection of the background monolayer revealed striped filamentous filaments forming spirals or concentric turns that often merged into wider sheets. Rings were usually observed in the spiral centre. After ∼15 minutes incubation, this dense mosaic of filaments spanned many microns in diameter (Figure 4D). By extracting and aligning 35 nm^2^ filament subsections, class averages were generated revealing a 54 Å spacing between filament stripes. Given this spacing and the width of the filaments was ∼12-14 nm, we concluded that these Vipp1 filaments were equivalent to those formed on SLBs. Overall, this data supported a model where spiral filaments and planar sheets have equivalent ultrastructure with the 54 Å repeat a key building block in polymer assembly.

**Figure 4.**
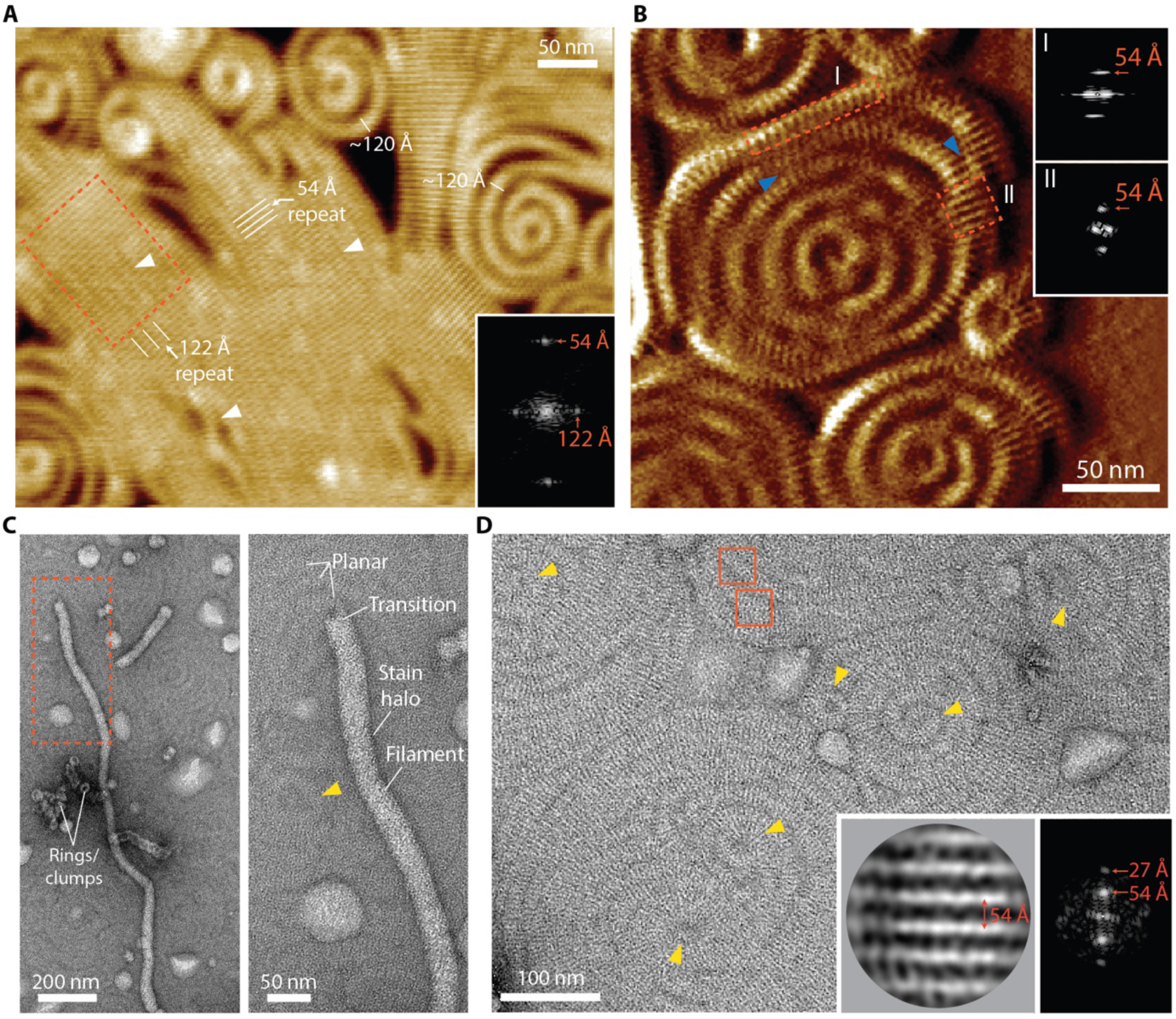
Vipp1 planar sheets and spiral filaments have closely related lattices. **A**, F-AFM phase image showing Vipp1 highly ordered planar sheets. Inset panel shows Fourier Transform of dotted box. Parallel ridges are spaced 54 Å apart. White arrows indicate merged filament seam lines with 122 Å repeat. **B**, F-AFM phase image showing Vipp1 spiral. Inset panels show Fourier Transform of dotted boxes. Parallel ridges are spaced 54 Å apart. Blue arrows show the merging of spiral turns into planar sheet showcasing their close polymeric relationship. **C**, NS EM image showing Vipp1 polymers (rings, rod-like filaments) decorating the surface of a monolayer which is itself covered by Vipp1 2D planar filaments. **D**, NS EM image showcasing the mosaic of 2D planar spirals and sheets Vipp1 forms on a lipid monolayer. Yellow arrows indicate rings at the centre of spirals. Red boxes indicate example regions for particle extraction and alignment. Inset shows particle class average (left) and corresponding Fourier Transform (right). Filament stripes are 54 Å apart.

### Vipp1Δα6_1-219_ truncation forms tightly packed planar spirals and highly ordered sheets

Truncation of the Vipp1 CTD (termed Vipp1Δα6_1-219_) somehow modulates Vipp1 polymerisation dynamics. Specifically, Vipp1Δα6_1-219_ purified in low salt conditions (10 mM NaCl) has a greater propensity to form ordered helical filaments and quasi-helical ring stacks (Supplementary Figure 3A) than Vipp1^1^. We therefore investigated how CTD removal would affect Vipp1 behaviour on SLBs and lipid monolayers. When mixed with SLBs and visualised by F-AFM, Vipp1Δα6_1-219_ bound the SLB edges like Vipp1. However, edges were coated with thin curled sheets or compact spirals where turns were usually interconnected with no central ring formed (Supplementary Figure 3B). Both spiral filaments and planar sheets had ∼6 nm height offsets from the membrane like Vipp1 (Supplementary Figure 3C). Remarkably, Vipp1Δα6_1-219_ had the capacity to form dense crystalline planar sheets up to a micron in width with parallel surface ridges 54 Å apart (Supplementary Figure 3D). Based on these dimensions, we concluded that Vipp1Δα6_1-219_ spirals and planar sheets have closely related ultrastructure to those of Vipp1. Overall, truncation of the CTD reduced filament dynamics so that spirals merged with a tendency to form sheets.

Importantly, detailed analysis of Vipp1Δα6_1-219_ sheets showed how they curl and gave important insights into rules that govern Vipp1 polymer curvature. When filaments curl, the rigid 54 Å-spaced substructure experiences tension or compression on the longer outside or shorter inside of the bend, respectively. Due to the width of sheets, curling occurs by adding filament sections into the outside of bends (wedging) whilst simultaneously removing sections on the inside (Supplementary Figure 3E and 3F). This exposes how flexibility between the 54 Å ridges is limited with only a defined squeezing or stretching of the spacing possible. This rule extends to the thinner spiral filaments, which are also governed by the 54 Å repeating substructure. Here, filament curvature is seldom induced by wedging (Figure 4B). Instead, filaments achieve high curvature either by inducing lattice breaks or rather by tilting and transitioning to 3D structures to alleviate elastic stress^40,47^.

Subsequent analysis of Vipp1Δα6_1-219_ exposed to lipid monolayer revealed rod-like filament structures bound to the surface that were transitioning to a planar-like array (Supplementary Figure 3G). In the background, a mosaic of planar filaments corralled around apparent raised regions and areas with ruptured monolayer (Supplementary Figure 3H). Consistent with the F-AFM data, filaments formed larger curved sheets so that distinct spirals were not readily observed (Supplementary Figure 3I). Notably, both a 54 Å and orthogonal 32 Å spacing was now detected, with the latter consistent with the axial rise between subunits in neighbouring rungs of Vipp1 rings^1^. Overall, our data supported a model where Vipp1Δα6_1-219_ builds planar polymers with the same substructure as Vipp1. The CTD has the effect of tuning Vipp1 polymer stability and dynamism. In its absence, Vipp1 has a higher avidity to form polymers with increased regularity and stability.

### Vipp1 helical filaments have a lattice closely related to Vipp1 rings_HS_

Given 2D planar sheets share a close geometric relationship with helical lattices^52^, and as a means of understanding the interconnectedness of Vipp1 polymers including rings, spirals and planar sheets, we determined the structure of four different Vipp1 helical filaments by cryo-EM (Table S1). Specifically, Vipp1 and a Vipp1 Interface 3 mutant F197K/L200K^1^ assembled using equivalent helical parameters and lattices (Supplementary Figure 4) and were termed Vipp1_L1_ and Vipp1_F197K/L200K_L1_, respectively. Alternatively, Vipp1Δα6_1-219_ assembled with related but different lattices termed Vipp1_Δα6_L2_ and Vipp1_Δα6_L3_. For all samples, conditions were screened to enrich for helical filaments although additional polymer types remained mixed. Vipp1Δα6_1-219_ formed longer and more stable helical filaments in comparison to native Vipp1^1^ with helical-like ribbons and rings observed (Figure 5A). In all samples, helical filaments were heterogeneous with multiple symmetries that required extensive *in-silico* classification to achieve near-uniform symmetry bins. Helical parameters could not initially be determined from low quality C1 symmetry reconstructions. Vipp1_Δα6_L3_ was the only lattice form where class averages of aligned particles yielded a Fourier Transform with non-overlapping layer lines amenable to indexing (Supplementary Figure 4). This gave a grid of possible symmetries which were systematically tested. Only helical parameters with rise = 2.16 Å and rotation = 85.50° gave a high-resolution map to 3.7 Å resolution (Figure 5B and Supplementary Figure 4A) with excellent side chain detail enabling model building from amino acids 1-217 (Figure 5C). Map quality was lowest around helix α5C indicative of instability within Interface 1 – a trait observed for all Vipp1 helical filaments to differing degrees. Vipp1_Δα6_L3_ was ∼24.4 nm in diameter with a hollow inner lumen diameter of ∼12.7 nm. Overall, the Vipp1 subunits self-assembled using similar interfaces as for Vipp1 ring_HS_ polymerisation^1^ indicating a close relationship between helical and ring polymers (Figure 5D-5F). Specifically, Vipp1 formed ESCRT-III-like protofilaments (Figure 5G-5I) where hairpin motifs packed side by side and the C-terminus of helix 5 (α5C) of subunit *j* bound across the hairpin tip of subunit *j*+3 to form the classical Interface 1 domain swop (Figure 5D and 5E). These protofilaments formed a 17-start right-handed helix that ran diagonally around the helical axis (Supplementary Figure 4B). Concurrently, each hairpin tip bound the N-terminus of helix 5 (α5N) in a neighbouring protofilament thereby forming Interface 3 (Figure 5D)^1^. Neighbouring protofilaments aligned laterally to form a 4-start left-handed helix where subunits connected through the stacking of helices α0 and the formation of Interface 2 (Figure 5F)^1^. Helix α0 lipid binding domains therefore lined and twisted around the inner lumen of the filament. The 4-start helix has a pitch of 44 Å and forms diagonal parallel ridges that encircle the helical axis and dominate the surface topology of the Vipp1 helical filament. For Vipp1_Δα6_L2_, the structure was closely related to Vipp1_Δα6_L3_ although different helical parameters yielded a modified helical lattice with a slight lattice rotation relative to the helical axis (Supplementary Figure 5A-C). Overall, the Vipp1_Δα6_L3_ and Vipp1_Δα6_L2_ structures revealed the close relationship between Vipp1 helical and ring_HS_ polymers highlighting how only relatively minor adjustments to assembly dynamics would facilitate transition between these forms.

**Figure 5.**
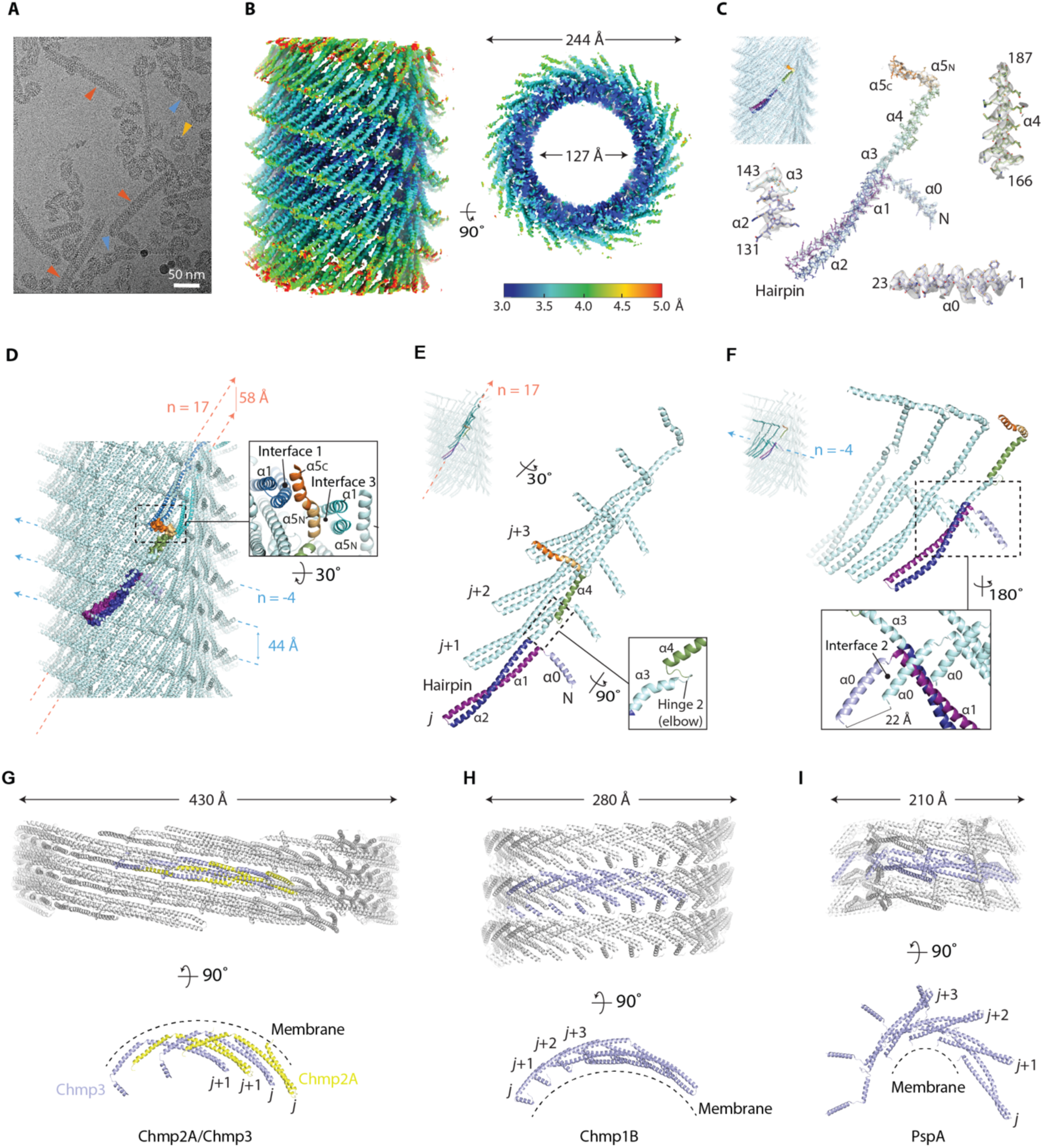
Vipp11′α6_1-219_ helical filaments have a lattice closely related to Vipp1 rings_HS_. **A**, Cryo-EM image showing Vipp11′α6_1-219_ forming helical filaments, helical-like ribbons, and rings (red, blue and yellow arrows, respectively). **B**, Sharpened Vipp1_Δα6_L3_ map contoured at 2.3α showing local resolution estimates. **C**, Vipp1_Δα6_L3_ map fitted with Vipp1_Δα6_L3_ helical filament structure (top left). The coloured monomer is isolated and zoomed to show map quality, build and fit. Map contoured at 3α except helix α5 at 1α. **D**, Structure of the Vipp1_Δα6_L3_ helical filament with zoom panel highlighting conservation of Interfaces 1 and 3. Bessel orders n=17 and n=-4 are indicated with pitch. **E**, The 17-start right-handed helix in Vipp1_Δα6_L3_ forms ESCRT-III-like protofilaments. **F**, The 4-start left-handed helix in Vipp1_Δα6_L3_ is formed by subunit stacking mediated by Interface 2. **G-I**, Helical structures of other ESCRT-III family members bound to membrane.

### Comparison of Vipp1 _L1_ with Vipp1Δα_6_L2_ and Vipp1Δα_6_L3_ revealed a mechanism for filament constriction

In comparison to Vipp11′α6_1-219_ sample, Vipp1 was characterised by unfurled ring stacks or helical-like ribbons with fewer and shorter ordered helical filaments suggesting a reduced stability for helical polymerisation (Supplementary Figure 6A). Although the Fourier transforms of Vipp1_L1_ and Vipp1_Δα6_L2_ class averages were similar with closely related lattices, only helical parameters with rise = 2.37 Å and rotation = -75.86° yielded a Vipp1_L1_ map to 3.7 Å resolution overall (Supplementary Figures 4 and 6B). Map local resolution range was broadest compared to other Vipp1 helical forms with inner parts of the filament well resolved compared to the periphery where the hairpin tips and helix α5 were more poorly ordered (Supplementary Figures 6C). The Vipp1_L1_ subunit was built between amino acids 1-214 (Supplementary Figures 6D and 6E) and assembled using similar interfaces as for Vipp1_Δα6_L2_ and Vipp1_Δα6_L3_. However, the Vipp1_L1_ subunit differed to Vipp1_Δα6_L2_ and Vipp1_Δα6_L3_ subunits with its hairpin tip kinked at Ile68 and helix α5 compacted (Supplementary Figures 6D and 6E) so that Interfaces 1 and 3 were perturbed. Although map potential relating to the CTD could not be assigned, our structures were consistent with this motif destabilising Interface 1 possibly through kinking of the hairpin whilst also directly or indirectly limiting Interface 3 formation. The effect of the CTD on Interfaces 1 and 3 thereby present a mechanism for Vipp1_L1_ helical filament instability and CTD-mediated tuning of Vipp1 polymer dynamics^1,30,31^. Compared with Vipp1_Δα6_L2_ and Vipp1_Δα6_L3_, the Vipp1_L1_ filament was constricted with a ∼21.0 nm and ∼10.5 nm external and inner diameter, respectively. The mechanism for this ∼2nm inner lumen constriction was through subunit removal. Specifically, the left-handed 21-start helix in Vipp1_Δα6_L2_ and Vipp1_Δα6_L3_ that runs almost parallel to the helix axis now formed a 19-start helix in Vipp1_L1_ (Supplementary Figure 4B). Concurrently, the number of ESCRT-III-like protofilaments was reduced to a 14-start right-handed helix. Filament constriction was mediated by angular adjustments of helices α3 and α4 resulting in a ∼7.5 Å upward flexing (Supplementary Figures 6E). These intra-subunit conformational changes were accompanied by subtle adjustments in subunit lattice packing resulting in a Vipp1_L1_ protofilament with higher curvature than Vipp1_Δα6_L2_ or Vipp1_Δα6_L3_ (Supplementary 6F and Video S4). Overall, the comparison of Vipp1_L1_, Vipp1_Δα6_L2_ and Vipp1_Δα6_L3_ filaments provided a mechanism for constriction where CTD-mediated adjustments to lattice assembly induced a reduction of ESCRT-III-like protofilaments and consequently a reduction in filament circumference. Our data was consistent with a model where the CTD played a key role in tuning Vipp1 assembly dynamics and has the capacity to drive helical filament constriction.

### Vipp1 polymers tubulate membrane along the same plane from different orientations

The Vipp1_F197K/L200K_ mutations impeded Interface 3 resulting in long helical filaments^1^ that often had a membrane vesicle cap at their tips (Figure 6A). Spheroid and helical tubule-like membrane vesicles coated in Vipp1_F197K/L200K_ were also observed. The membrane was presumably bound during purification from *E. coli*. Vipp1_F197K/L200K_L1_ was resolved to 3.7 Å overall (Figure 6B and Supplementary Figure 4A) using helical parameters of rise = 2.44 Å and rotation = -75.83°, which were almost identical to those of Vipp1_L1_. Excellent map quality and side chain detail facilitated model building (Supplementary Figure 5D). Notably, this included flexible hinge 2, which was well ordered in comparison to other Vipp1 maps. Overall, Vipp1_F197K/L200K_L1_ and Vipp1_L1_ subunits were in similar conformations (Supplementary Figure 5E) and assembled using equivalent helical lattices (Figure 6C and Supplementary Figure 4). However, due to the Vipp1_F197K/L200K_ mutation, Interface 3 formation was inhibited with no supporting map and build for helix α5N. No map potential was located for C-terminal helix α6. The key difference between Vipp1_F197K/L200K_L1_ and Vipp1_L1_ maps was the presence of lipid bilayer in the central lumen. Here, the inner leaflet formed a 4 nm diameter tube, which is close to the limit where hemifusion is expected to occur, whilst the outer leaflet filled the space between the helix α0 stacks (Figure 6D). Analysis of the Vipp1_F197K/L200K_L1_ electrostatic surface potential showed amphipathic helix α0 to be positively charged with basic residues positioned to attract negatively charged lipid headgroups. Hydrophobic residues oriented along the inner lumen face then contact the fatty acid chains (Figure 6E). Importantly, the structure of Vipp1_F197K/L200K_L1_ facilitated a comparison of Vipp1 helical and ring_HS_ polymers when bound to membrane. In the C14 symmetry Vipp1 ring_HS_, the ESCRT-III-like protofilaments formed rungs that were orthogonal to the membrane bud/tube axis^1^. In Vipp1_F197K/L200K_L1_, the ESCRT-III-like protofilaments were rotated ∼43° to the membrane tube axis whilst maintaining similar lateral interactions and ultrastructure by undergoing filament twist (Figure 6F). Therefore, by forming a 3D pliable polymer, the Vipp1 lattice can induce membrane tubulation along the same axis but by using radically different lattice orientations. To facilitate the transition from Vipp1 ring_HS_ to helical polymer, significant conformational changes were observed within individual subunits to accommodate the relative shift in membrane plane (Figure 6G and Video S5). Mediated by Hinges 1 and 2, helices α3 and α4 flexed upwards and outwards ∼9° and ∼20°, respectively. Concurrently, Hinge 3 facilitated additional shifts in helix α5C orientation cumulating in a ∼52 Å swing as measured for Ala211 in helix α5C. In addition, the length of the Vipp1 subunit was reduced from 158 Å in Vipp1 ring_HS_ to 149 Å in Vipp1_F197K/L200K_L1_ between the hairpin tip and helix α4 terminus. This important compaction was mediated by Hinge 1 with the N-terminus of helix α4 now overlapping the C-terminus of helix α3. Overall, Vipp1_F197K/L200K_L1_ showed how *N. punctiforme* Vipp1 helical filaments have the capacity to tubulate membrane, as reported in other Vipp1 systems^53^. Moreover, it shows that the transition between helical and ring conformations relative to the membrane plane is induced by lattice rotation and filament twist rather than through a novel polymer form. This is important as the transition of ESCRT-III planar spirals and rings to 3D spirals likely requires filament rotation relative to the membrane plane, as has already been glimpsed^26,33,50^.

**Figure 6.**
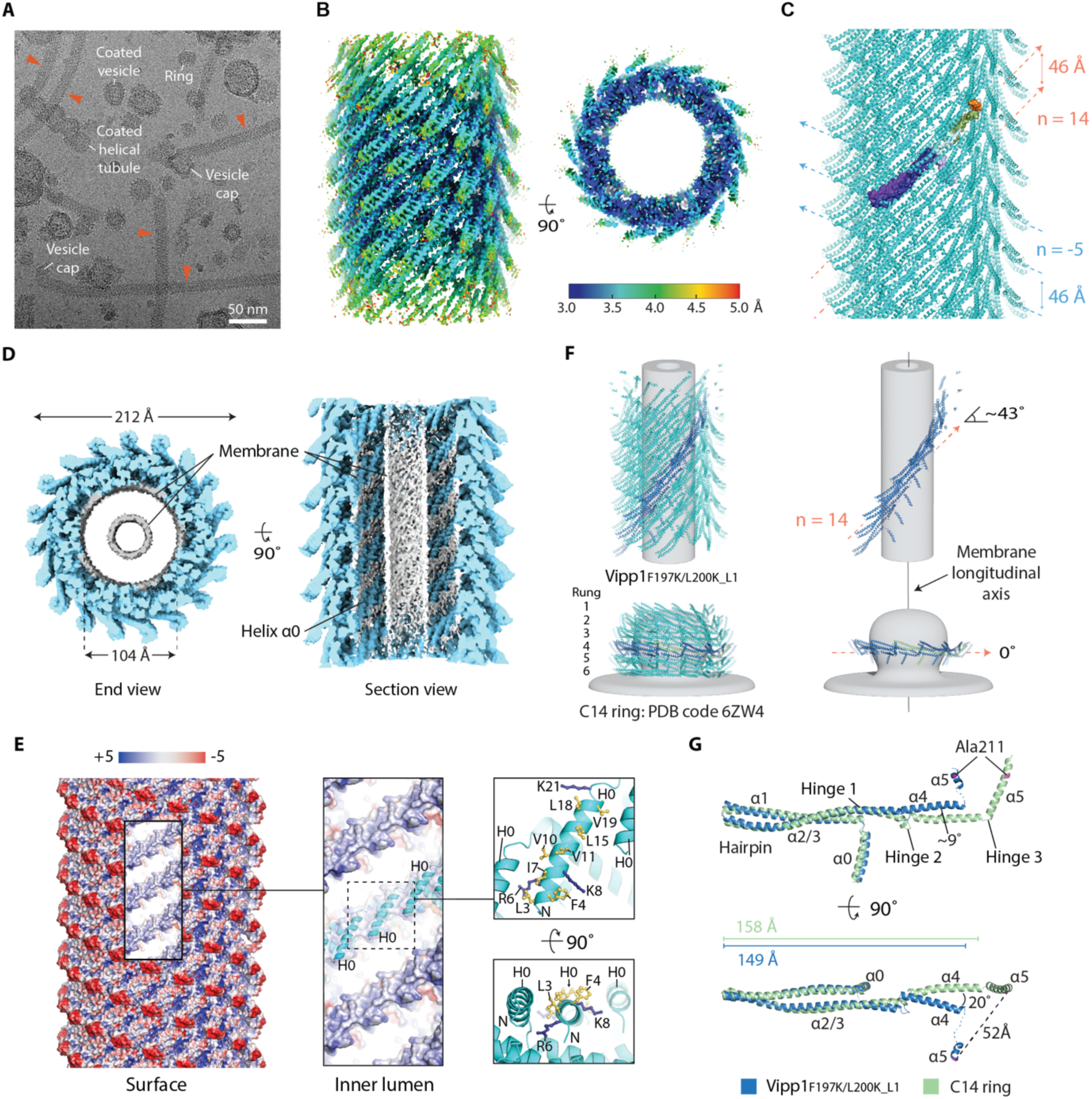
Vipp1_F197K/L200K_L1_ is constricted and tubulates membrane. **A**, Cryo-EM image showing Vipp1_F197K/L200K_L1_ forming helical filaments and coated membrane tubules. **B**, Sharpened Vipp1_F197K/L200K_L1_ map contoured at 4α showing local resolution estimates. **C**, Structure of the Vipp1_F197K/L200K_L1_ helical filament with one monomer coloured. Bessel orders n=14 and n=-5 are indicated with pitch. **D**, Unsharpened Vipp1_F197K/L200K_L1_ map contoured at 3α showcasing tubulated membrane within the inner lumen. **E**, Vipp1_F197K/L200K_L1_ filament surface rendered to show electrostatic charge. Blue to red spectrum represent positive to negative charges with units *k_B_T*/*e_c_*. Zoom panels show mechanism of membrane binding. **F**, Comparison of ESCRT-III-like protofilament orientation relative to the same membrane plane between Vipp1_F197K/L200K_L1_ and Vipp1 ring_HS_. **G**, Hairpin superposition of a Vipp1_F197K/L200K_L1_ subunit with a Vipp1 ring_HS_ subunit (C14 symmetry, rung 4, PDB code 6ZW4).

## Discussion

Here we show how the bacterial ESCRT-III-like protein Vipp1 assembles dynamic planar spirals and sheets on membrane supports. Moreover, the spirals assemble central rings that protrude and abscise from the parent filament. These rings share dimensions with Vipp1_HS_ rings known to bud membrane. In addition, we show the structure of four different Vipp1 helical filaments with three related but different lattice assemblies. They show the close connection between 2D planar, helical and Vipp1_HR_ ring polymers with the transition between the latter two achieved by a remarkable lattice rotation and twist (Figure 7A) around the same membrane plane. Given these different polymer forms are highly ordered, their comparison provides a basis for modelling how Vipp1 builds planar filaments that transition to 3D membrane budding forms. Collectively, our results suggest a general mechanism for how Vipp1 may stabilise and repair membrane.

**Figure 7.**
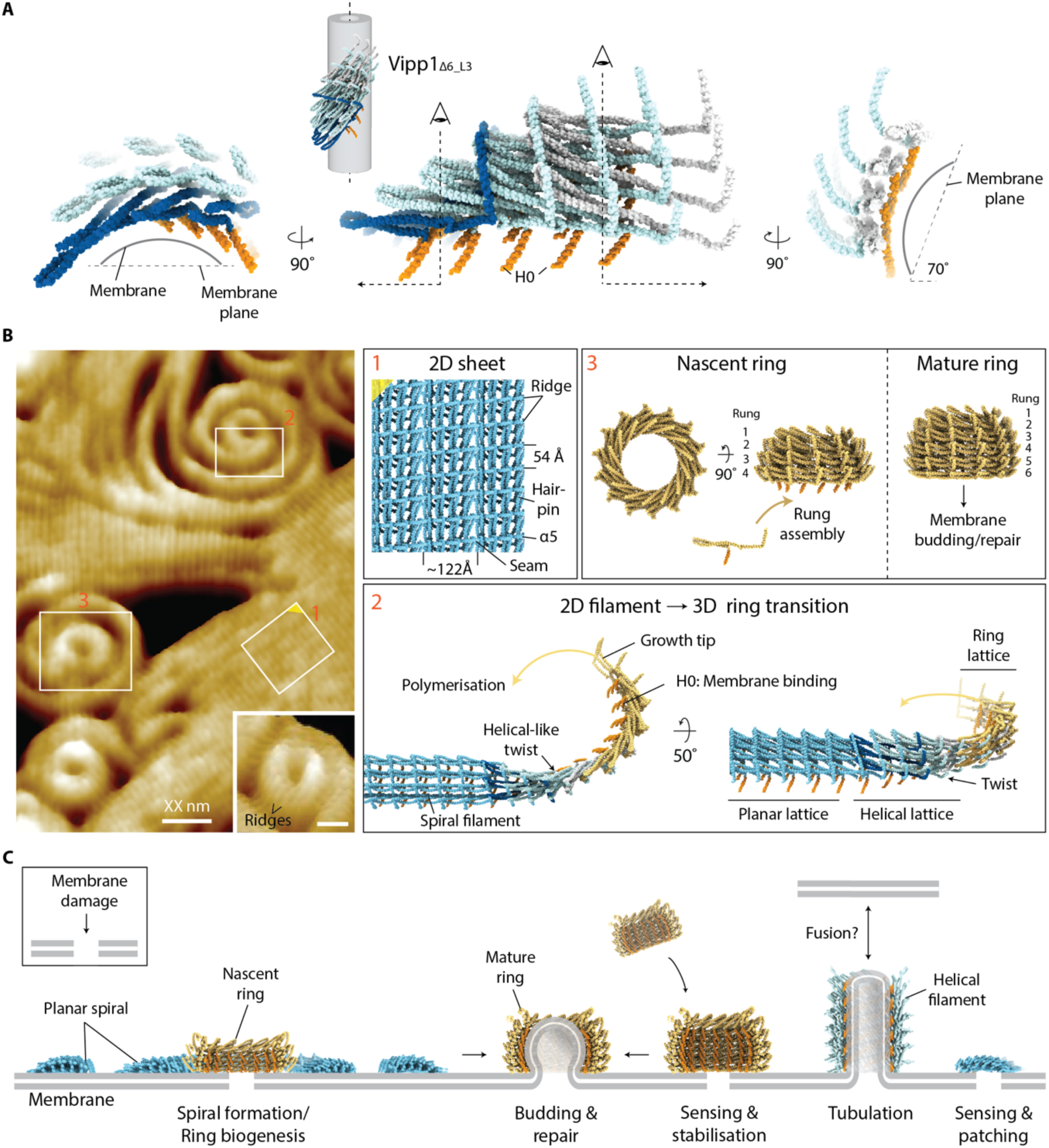
Mechanism for Vipp1 spiral formation, ring biogenesis and membrane repair. **A**, Section of four ESCRT-III-like protofilaments extracted from the Vipp1_1′α6_L3_ helical structure showing how filament twist enables binding of membranes on opposing planes. **B**, Model of Vipp1 planar sheet, spiral and 3D ring biogenesis. F-AFM image (left) with white boxes 1-3 relating to right panels 1-3. Note how the twisting helical filament in **A** bridges the transition from planar to 3D ring structures. To facilitate modelling of planar filament helix α4 was removed. **C**, Mechanism of Vipp1-mediated membrane sensing, stabilisation and repair. In *Chlamydomonas*, apparent Vipp1 helical filaments bridge thylakoids and the chloroplast envelope ^34^.

Both F-AFM and NS-EM data show Vipp1 planar sheets and filaments to be characterised by parallel ridges with a 54 Å repeat (Figure 4A, 4B, and 4D). Additionally, the Vipp11′α6_1-219_ planar filaments include a 32 Å repeat orthogonal to the 54 Å spacing (Supplementary Figure 3I). By calculating the cylindrical projection^54^ of the Vipp1_Δα6_L3_ helical filament (Supplementary Figure 7A) the 3D map may be represented as a geometrically equivalent 2D lattice. The 4-start left-handed helix is the dominant feature forming parallel stripes with 44 Å pitch. These stripes relate to the ridges on the Vipp1_Δα6_L3_ filament surface when visualised by cryogenic electron tomography (Supplementary Figure 7B). Inter-ridge distance is formed by the spacing between neighbouring hairpins within each ESCRT-III-like filament (Supplementary Figure 7C). Intriguingly, inter-hairpin distance has a degree of flexibility with hairpins sliding up to 14 Å relative to each other in Vipp1_HS_ rings depending on rung position (Supplementary Figure 7D). In Vipp1_HS_ C17 symmetry rings, hairpin spacing spans 53-59 Å in the central rungs with each neighbouring ESCRT-III-like filament 32-33 Å apart (Supplementary Figure 7C). Overall, the lattice dimensions of helical and ring_HS_ polymers are consistent with those obtained from the Fourier Transform of Vipp1 and Vipp11′α6_1-219_ planar sheets and filaments on a lipid monolayer (Figure 4D and Supplementary Figure 3I) and SLBs (Figure 4A and 4B). Our data therefore supports a model where Vipp1 planar sheets and spiral filaments are geometrically similar to unfurled and flattened Vipp1 rings_HS_ or helical filaments with the key assembly Interfaces 1-3 maintained. Sheets and spiral filaments comprise parallel ESCRT-III-like protofilaments (Figure 5E) with the ridges running near orthogonal to the protofilament axis (Figure 7B). The 122 Å spacing observed as seams within the sheets is close to the F-AFM-measured ∼13 nm width of spiral filaments (Figure 2G) and is consistent with spiral filaments comprising four or sometimes five parallel protofilaments. Snf7 spirals generally form from just one protofilament, which likely explains their increased compressibility and deformation into polygons^40^.

As Vipp1 spiral filaments grow centrally, curvature increases with each turn building elastic stress^40,47^. For spiral filaments to curve, our data supports a model where hairpin sliding and subunit flexing via hinges 1-3 enable each planar filament to curl laterally on the membrane plane with the inter-ridge distance shortening on the inside of the filament and extending on the outside. However, based on measurements from different rungs in Vipp1 rings_HS_^1^, inter-ridge distance cannot compress below ∼41 Å or stretch greater than ∼61 Å (Supplementary Figure 7D). These distances are insufficient to facilitate the high level of curvature observed in the filaments or rings_LS_ at the centre of spirals if in a strictly planar form. Therefore, once inter-ridge distance limits are reached, residual curvature-induced elastic stress must now be minimised either by breaking the filament, wedging (Supplementary Figure 3F) or through filament tilt and transitioning to a 3D form. For Vipp1 filaments, tilt is observed by mounting height offset from the membrane particularly within central turns of the spiral (Figure 3C and Supplementary Figure 1A).

In the centre of each Vipp1 spiral where filament curvature was highest, Vipp1 rings_LS_ formed which protruded an additional ∼1 nm above the surrounding spiral filaments due to filament tilt. Ridges were sometimes observable on the outside face of Vipp1 rings_LS_ showcasing filament tilt (Figure 7B). Importantly, the filament twist observed in Vipp1 helical polymers (Figure 7A) provided a mechanism for how ESCRT-III-like filaments tilt and bind membrane on different planes thereby enabling Vipp1 to transition from planar to 3D ring architectures (Figure 7B). In ESCRT-III, changes in geometry of membrane bound filaments, including filament tilt, underlie membrane deformation from a planar spiral to a 3D helix ^26,48,49^. Ultimately, filament tilt in Vipp1 rings_LS_ relative to the parent spiral induce torsion and promote abscission. Although Vipp1 rings_HS_ have a substantially lower height when measured by F-AFM (Supplementary Figure 2C) in comparison to structural measurements^1^, our data was consistent with Vipp1 rings_LS_ constituting ∼3-4 rungs rather than 5-7 rungs in a mature Vipp1 ring_HS_ (Figure 7B). Final maturation steps may require a non-rigid membrane support, changes in membrane composition or additional factors such as nucleotide^34,55^.

Collectively, our results provide a mechanism for how Vipp1 functions in membrane stabilisation and repair (Figure 7C) in cyanobacteria and chloroplasts. Both Vipp1 small oligomers (or monomers) and rings_HS_ sense and bind highly curved and perturbed membrane. This finding is consistent with Vipp1 locating to the highly curved edges of thylakoid membranes^18^. Depending on the conditions, the small oligomers can polymerise into spirals that encircle the damaged region, which may incorporate protein complexes^19,56,57^, ultimately assembling a central ring structure. Mature Vipp1 rings_HS_ have the capacity to bud membrane thereby supporting a repair mechanism based on membrane squeezing and fusion^1^. Alternatively, by tuning the CTD, spiral filaments can readily anneal to form planar crystalline concentric or linear sheets (Supplementary Figure 3D), which would stabilise damaged membrane acting as a physical barrier and supporting scaffold. Such structures explain how Vipp1 and other bacterial homologues such as PspA might inhibit proton leakage across membranes^46,58^. Finally, Vipp1 helical filaments may form membrane bridges linking thylakoids and the chloroplast envelope^34^.

In conclusion, our results utilise Vipp1 as a model system to show how planar filaments transition to 3D rings. The structural homology between Vipp1 and other ESCRT-III proteins suggest that the basic principles observed here, including conserved ultrastructure between planar and 3D forms, filament tilt and twist, and lattice rotation on an equivalent membrane plane, will extend to other family members.

## Acknowledgements

Cryo-EM data was collected at Diamond Light Source and London Consortium for CryoEM (LonCEM). We thank Nora Cronin (LonCEM, The Francis Crick Institute) for cryo-EM data collection support, Paul Simpson for in-house EM support, and Suhail Islam for in-house computational support. We thank Diamond for access and support of the cryo-EM facilities at the UK national electron bio-imaging centre (eBIC) funded by the Wellcome Trust, MRC and BBSRC. JS was funded by a BBSRC grant (BB/W008181/1) to HL. This work was funded by a Wellcome Trust Senior Research Fellowship (215553/Z/19/Z) to HL. AR acknowledges funding from the Swiss National Fund for Research, grants N°310030_200793/1 and N°CRSII5_189996, and from the European Research Council, Synergy Grant N°951324_R2-TENSION. JE acknowledges support from EMBO long-term post-doctoral fellowship ALTF 989-2022. AC acknowledges funding from MCIU, PID2022-140687NB-I00; MCIU/AEI/FEDER MINECOG19/P66, RYC2018-024686-I, and Basque Government IT1625-22.

## Author contributions

JE and AR conceived the in-vitro assays for photonic microscopy. JE performed all photonic microscopy experiments. AM and AC designed and conceived Fast-AFM experiments. AM and AC gathered and analysed Fast-AFM data. SN and HL designed and conceived the EM studies. SN purified Vipp1 samples and undertook NS- and cryo-EM studies. SN and HL collected and processed cryo-EM data and built helical structures. JS contributed to Vipp1 sample purifications. AC and HL analysed the data and wrote the paper with contributions and revisions from all authors.

---

**Supplementary Figure 1.**
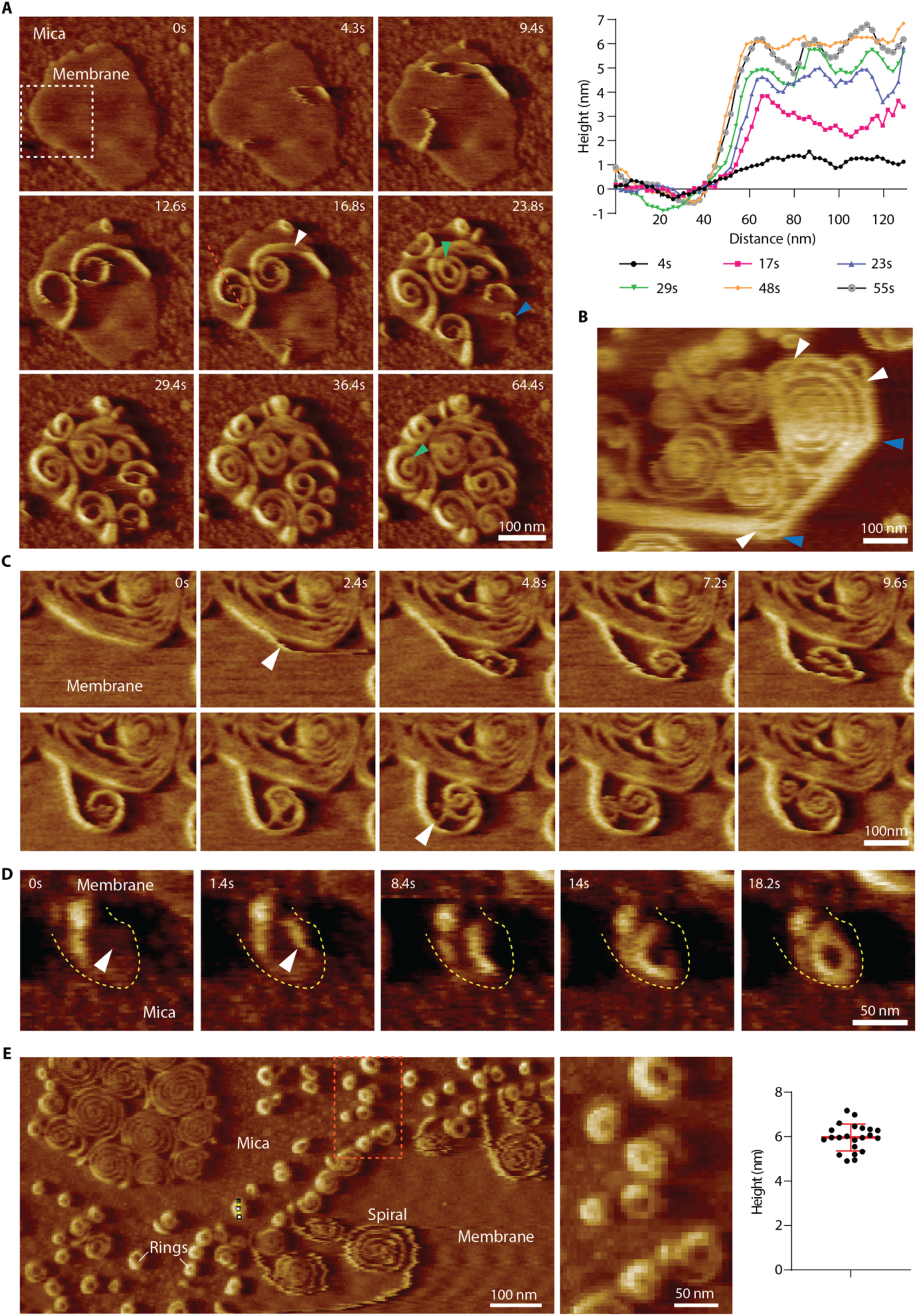
Vipp1 assembles dynamic networks of sheets, spirals and rings on membrane, related to Figures 2 and 3. **A**, Phase F-AFM timecourse showing spiral formation and ring_LS_ biogenesis. Highlighted events include ring_LS_ abscission, filament branching, and ring formation in the absence of a spiral (green, white and blue arrows, respectively). White dotted box indicates zoomed region relating to Figure 3A. Red dotted line relates to plotted height profile (right). **B**, Phase F-AFM image showing the outer turn of a spiral formed by linear sheet connected by vertices. Here, lattice breaks in the filament packing marked shifts in polymerisation direction (blue arrows). White arrow indicates sheet branching. **C**, Phase F-AFM timecourse showing how filaments often grow alongside established filaments with a propensity to branch off and form spirals (white arrows). **D**, Phase F-AFM timecourse showing ring formation due to spatial constraint (white arrow). Dotted yellow line delineates membrane boundary. **E**, Phase F-AFM image showing examples of rings formed in the absence of a spiral. These rings form on membrane micro-patches on the mica due to spatial constraint. Middle panel relates to dotted box in left panel. The plot (right) shows the rings with mean height 6.0 ± 0.6 nm, N=22.

**Supplementary Figure 2.**
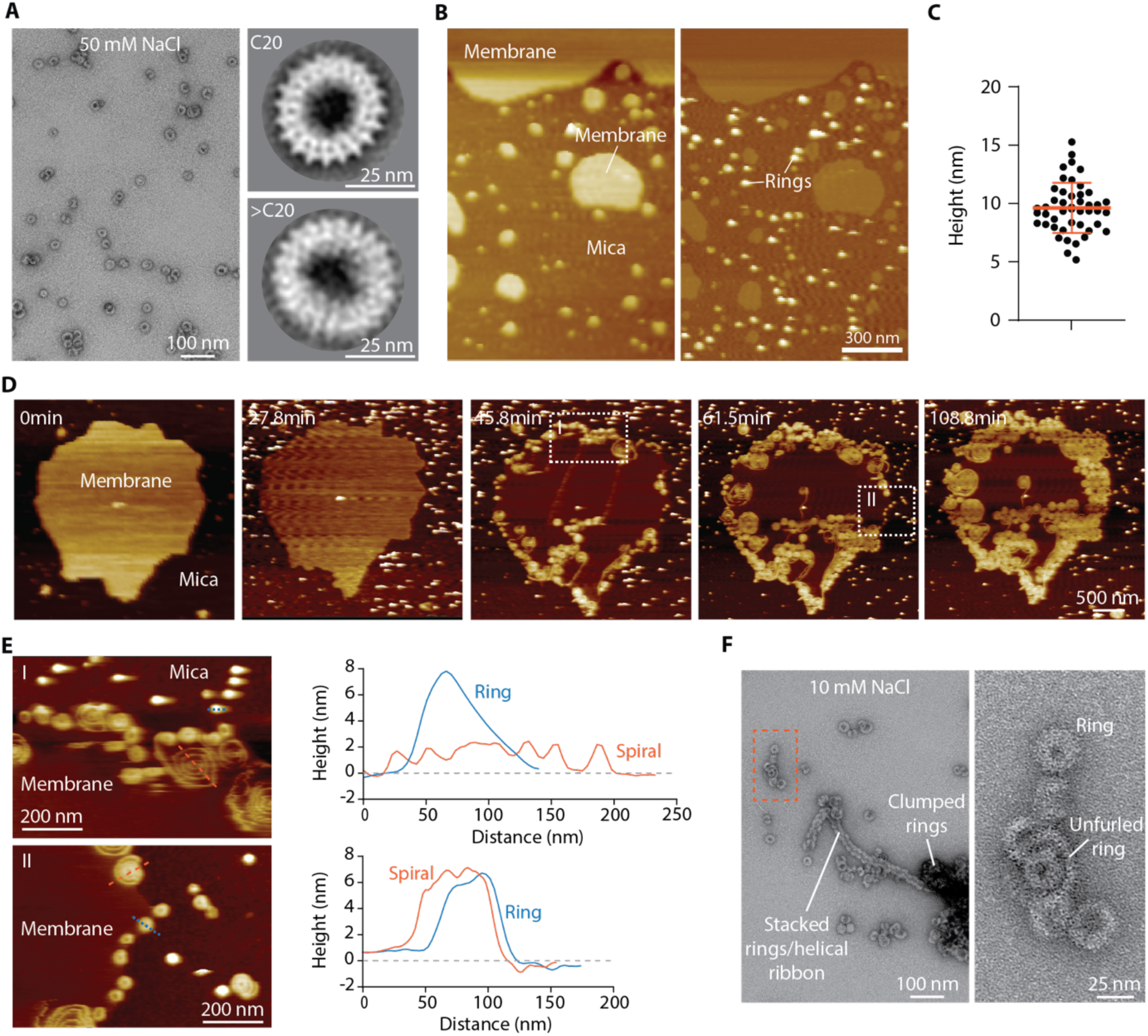
Vipp1 rings_HS_ form planar spirals in low salt conditions, related to Figure 3. **A**, Vipp1 rings_HS_ purified in high salt (50 mM NaCl) form rings between C11-C17 symmetries^1^ but also with higher symmetries such as C20 or above with diameters of 41 nm and 43 nm, respectively. **B**, Vipp1 rings_HS_ bind to mica as stable pre-formed dome-shaped rings^1^ in a high salt (50 mM NaCl) buffer. This panel is complementary to Figure 3H. **C**, Plot showing Vipp1 rings_HS_ heights as measured by F-AFM. Mean height is lower than Vipp1 rings_HS_ heights as determined by cryo-EM between ∼15-21 nm for C11-C17 symmetries^1^. **D**, F-AFM phase timecourse showing how, in a low salt (10 mM NaCl) buffer, Vipp1 rings_HS_ sample forms planar sheets and spiral networks on membrane. **E**, F-AFM image highlighting dotted box regions I and II in **D**. Rings and spirals nucleate and assemble on the highly curved membrane edge. Blue and red dotted lines were used for plotted height profiles (right) of typical spiral and rings. **F**, NS-EM image of Vipp1 rings_HS_ sample after buffer exchange into 10 mM NaCl buffer. Compared with the sample purified in 50 mM NaCl as in **A**, rings tend to be clumped or stacked and in the process of unfurling. Dotted box indicates zoomed region (right panel).

**Supplementary Figure 3.**
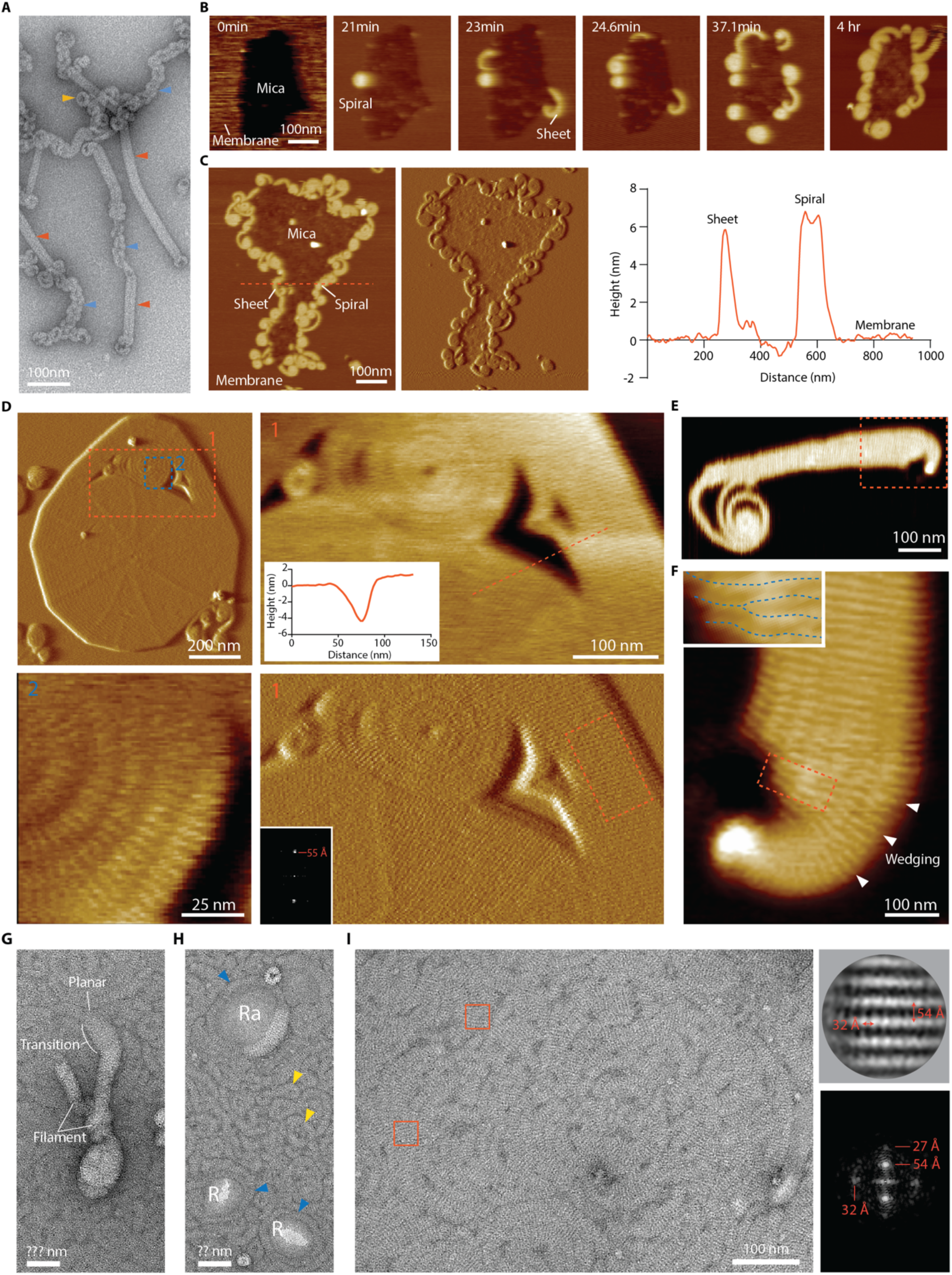
Vipp11′α6_1-219_ truncation forms tightly packed planar spirals and highly ordered sheets, related to Figures 2-4. **A**, NS-EM image showing Vipp11′α6_1-219_ forming helical filaments, helical-like ribbons, and rings (red, blue and yellow arrows, respectively). Related to Figure 5A. **B**, F-AFM phase timecourse showing Vipp11′α6_1-219_ recruitment to the membrane where it forms curled sheets and compact spirals on the membrane edge. **C**, F-AFM height and phase image (left) showing membrane edges fully decorated with sheet and compact spirals. Red dotted line indicates plotted height profiles (right) with Vipp11′α6_1-219_ planar filaments an equivalent height to native Vipp1, related to Figure 2E. **D**, Vipp11′α6_1-219_ form dense crystalline planar sheets and spirals. Dotted red line in top right panel was used for the plotted height profile (inset). Dotted box in bottom left panel indicates region used for Fourier Transform (inset). Parallel ridges are separated by a 55 Å spacing. **E**, F-AFM phase image showing Vipp11′α6_1-219_ planar sheet curling at one end and branching into a spiral at the other. **F**, Zoom of dotted box in **E**, showing how planar sheets curl via a wedging mechanism on the outer side of the bend (white arrows). Concurrently, sections are removed from the inner side (red dotted box and inset). **G**, NS EM image showing Vipp11′α6_1-219_ rod-like filaments decorating the surface of a monolayer which is itself covered by 2D planar filaments. **H**, Vipp11′α6_1-219_ forms a mosaic of 2D planar filaments (blue arrows) and rings (yellow arrows) on a lipid monolayer. Filaments encircle apparent raised areas (Ra) and regions where the monolayer is ruptured (R). **I**, Zoom view of Vipp11′α6_1-219_ mosaic of 2D planar filaments and sheets on a lipid monolayer. Note spiral filaments are not observed here, related to Figure 4D. Red boxes indicate example regions for particle extraction and alignment. Top right shows particle class average and corresponding Fourier Transform (bottom right). Filament stripes are 54 Å apart with an orthogonal 32 Å repeat.

**Supplementary Figure 4.**
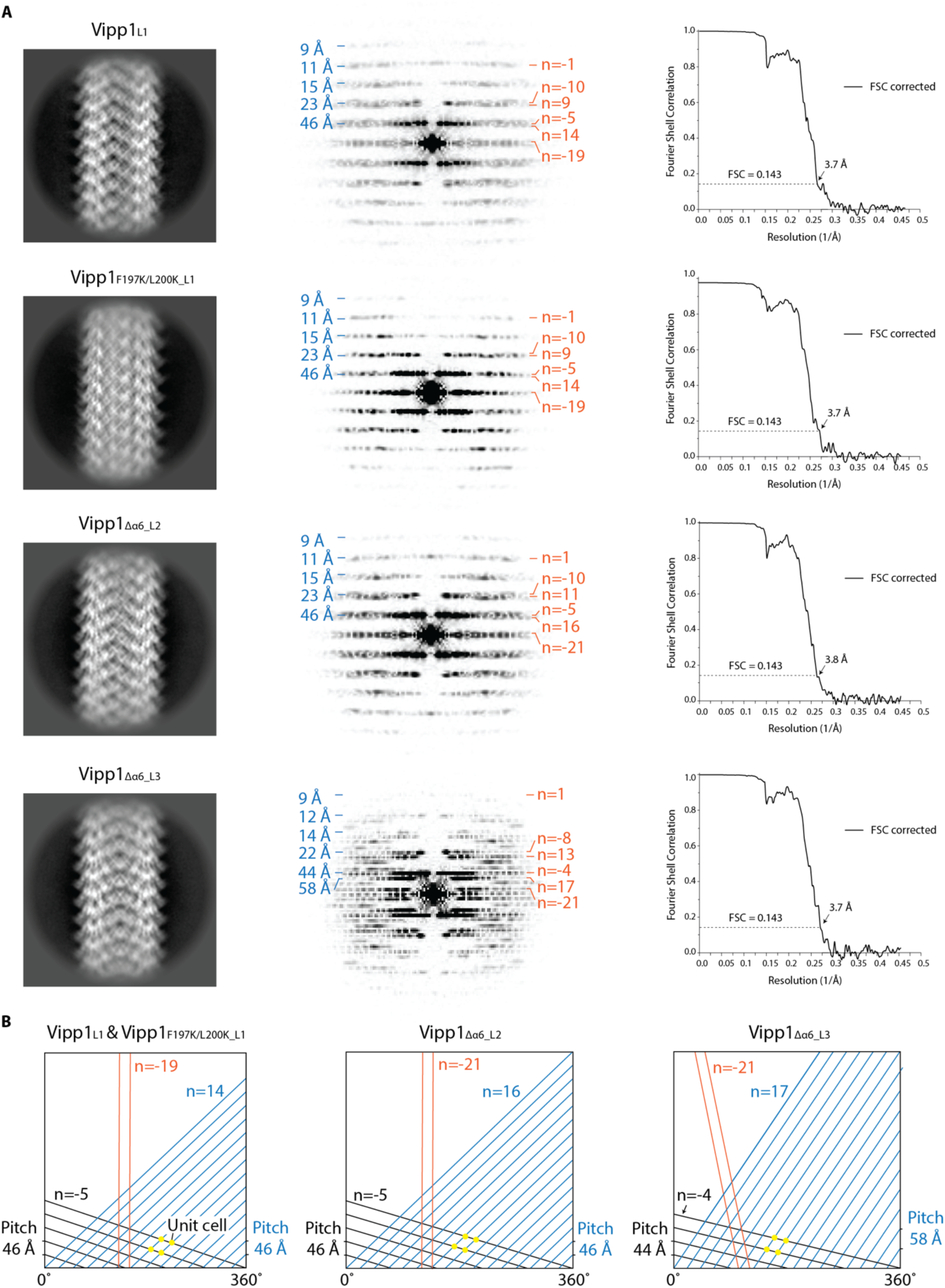
Vipp1 helical lattice analyses and map resolutions. **A**, 2D class averages of four different helical filaments including Vipp1_L1_, Vipp1_F197K/L200K_L1_, Vipp1_1′α6_L2_ and Vipp1_1′α6_L3_ (left column) with Fourier Transforms and Bessel Orders for key layer lines assigned (middle columns). Associated gold standard FSC curves for each map are presented (right column). **B**, Vipp1_L1_, Vipp1_F197K/L200K_L1_, Vipp1_1′α6_L2_ and Vipp1_1′α6_L3_ helical assemblies drawn as 2D lattices showing arrangement of key Bessel orders.

**Supplementary Figure 5.**
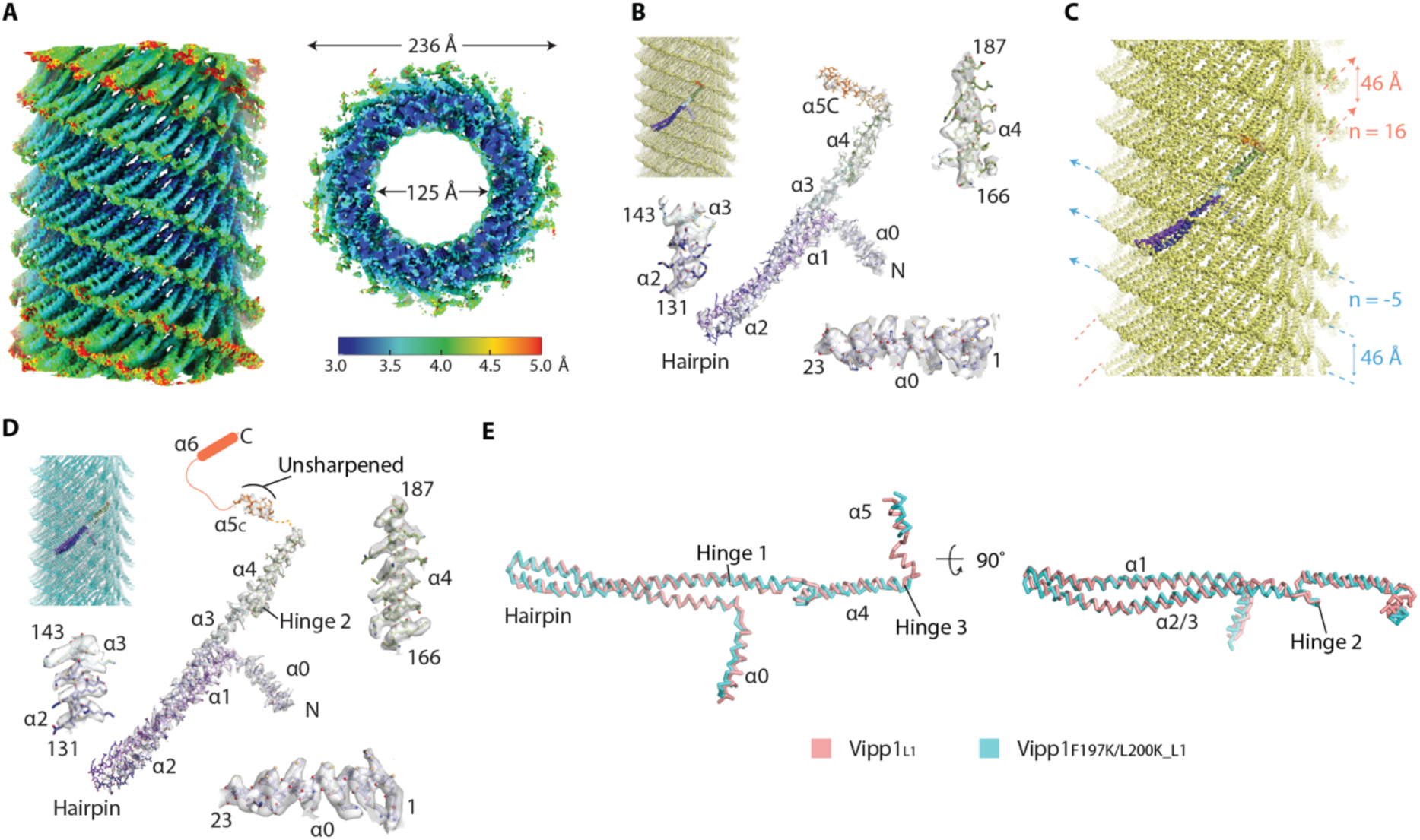
Cryo-EM maps and model build for Vipp11′α_6_L2_ and Vipp1_F197K/L200K_L1_, related to Figures 5 and 6. **A**, Sharpened Vipp1_1′α6_L2_ map contoured at 2σ showing local resolution estimates. Vipp1_1′α6_L2_ map was resolved to 3.8 Å overall using helical parameters rise = 2.16 Å and rotation = 68.50°. **B**, Vipp1_1′α6_L2_ map fitted with Vipp1_1′α6_L2_ helical filament structure (top left). The coloured monomer is isolated and zoomed to show map quality, build and fit. Map contoured between 3-4σ. Map local resolution range was broader than for Vipp1_1′α6_L3_ with generally lower map quality around helix α5. The Vipp1_1′α6_L2_ lattice therefore has increased instability in comparison to Vipp1_1′α6_L3_ although residual heterogeneity in the particle stack cannot be discounted as a contributory factor in reducing map quality. **C**, Structure of the Vipp1_1′α6_L2_ helical filament. Bessel orders n=16 and n=-5 are indicated with pitch. Specifically, the ESCRT-III-like protofilaments form a 16-start right-handed helix which twist around the helical axis with a 46 Å pitch. Concurrently, these protofilaments align to form a 5-start left-handed helix with a 46 Å pitch. In comparison to Vipp1_1′α6_L3_ (Figure 5 and Supplementary Figure 4), the effect is to rotate the Vipp1 lattice ∼13° clockwise around the helical axis with modest packing adjustments accommodating a minor flattening of the rhomboid unit cell. This results in a slightly narrower filament ∼23.6 nm in diameter with a hollow inner lumen diameter of ∼12.5 nm. Overall, the Vipp1_1′α6_L2_ and Vipp1_1′α6_L3_ structures reveal how subtle shifts in Vipp1 assembly and lattice packing modulate inherent polymer stability. **D**, Vipp1_F197K/L200K_L1_ map fitted with Vipp1_F197K/L200K_L1_ helical filament structure (top left). The coloured monomer is isolated and zoomed to show map quality, build and fit. Map contoured at 4σ. Related to Figure 6. **E**, Superposition of subunits from Vipp1_F197K/L200K_L1_ and Vipp1_L1_ with RMSD Cα = 1.8 Å. Subunits share similar conformations although Vipp1_F197K/L200K_L1_ does not have the hairpin kink observed in Vipp1_L1_ (Supplementary Figure 6D).

**Supplementary Figure 6.**
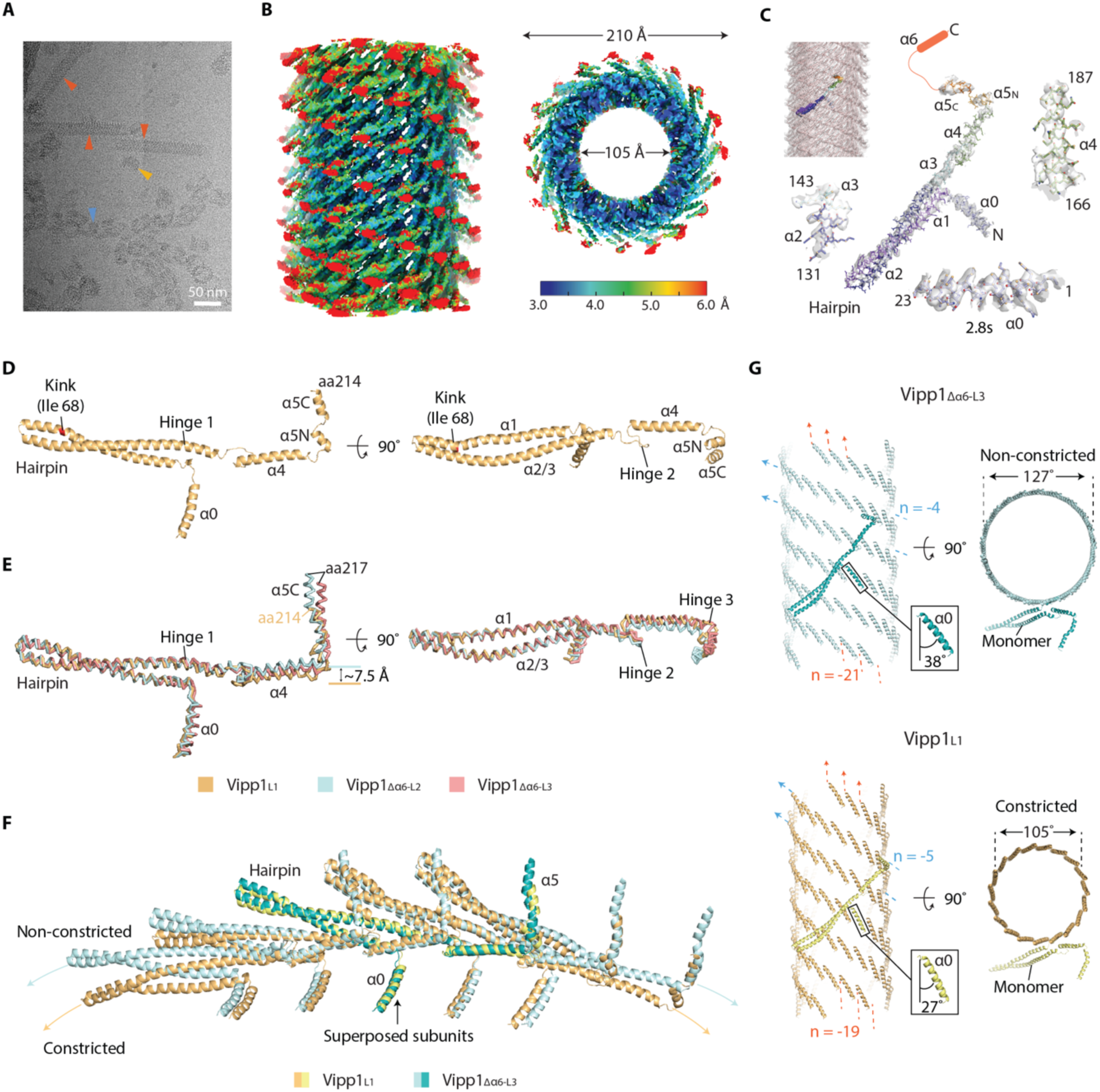
Comparison of Vipp1_L1_ with Vipp11′α_6_L3_ yields a mechanism of filament constriction. **A**, Cryo-EM image showing Vipp1_L1_ forming helical filaments, helical-like ribbons, and rings (red, blue and yellow arrows, respectively). **B**, DeepEMhancer^59^ post-processed Vipp1_L1_ map contoured at 3σ showing local resolution estimates. **C**, Vipp1_L1_ map fitted with Vipp1_L1_ helical filament structure (top left). The coloured monomer is isolated and zoomed to show map quality, build and fit. Map contoured between 2.8-3.5σ except helix α5c which was not post-processed. **D**, Structure of a Vipp1_L1_ subunit. **E**, Superposition of Vipp1_L1_, Vipp1_1′α6_L2_ and Vipp1_1′α6_L3_ monomers with the alignment focussed on the hairpin motif. The Cα RMSD of Vipp1_L1_ subunits to Vipp1_1′α6_L2_ or Vipp1_1′α6_L3_ was 1.5 Å and 1.7 Å, respectively. Note how helix α5 is compacted in Vipp1_L1_ compared with Vipp1_1′α6_L2_ and Vipp1_1′α6_L3_. **F**, Comparison of constricted versus non-constricted ESCRT-III-like protofilaments in Vipp1_L1_ and Vipp1_1′α6_L3_. Five monomers were superposed with the alignment focussed on the central subunit. **G**, Helical constriction occurs through subunit loss from the circumferential belt. Vipp1_1′α6_L3_ and Vipp1_L1_ filaments are compared, with constriction occurring due to a loss of two subunits (n= -21 reduced to n= -19) coupled with ∼11 Å rotation of the left-handed four-start helix as indicated in the helix α0 zoom boxes.

**Supplementary Figure 7.**
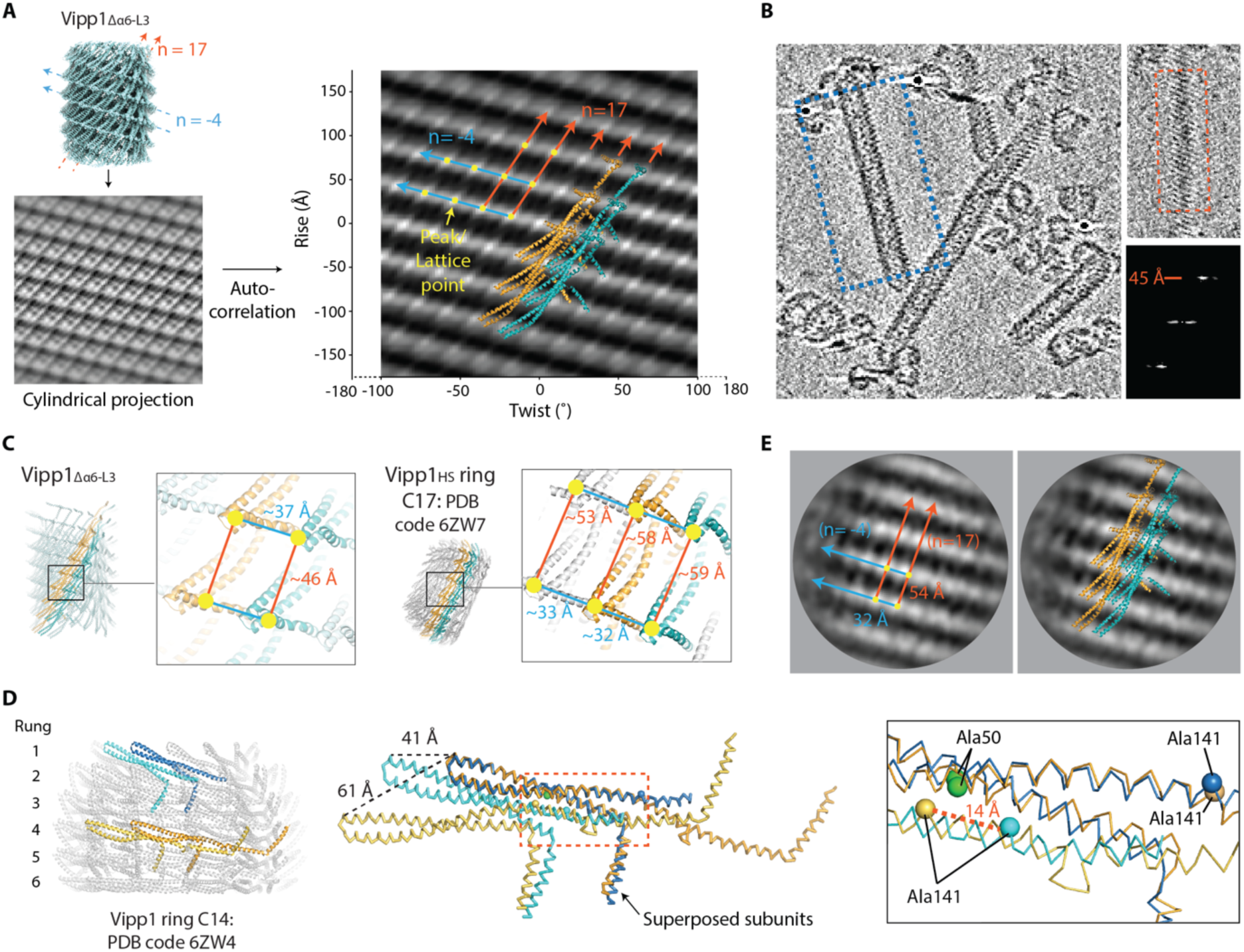
Analysis of Vipp1 planar sheet and spiral ultrastructure. **A**, Cylindrical projection of the Vipp1_1′α6_L3_ map with its auto-correlation function (Sun et al. 2022). Based on the Vipp1_1′α6_L3_ structure, the orientation of six subunits have been superposed on the 2D lattice with the ESCRT-III-like protofilaments coloured orange and turquoise. **B**, Cryogenic electron tomogram of Vipp1_1′α6_L3_. Central slice through the helical filaments is shown (left panel). Top right panel shows blue dotted box region from left panel. Only the upper surface ridges or stripes of the helical filament are sliced here. Bottom right panel shows Fourier Transform of red dotted box region. The ridges have a 45 Å repeat, which is consistent with the Vipp1_1′α6_L3_ left-handed 4-start helix pitch. **C**, Inter-subunit packing measurements of Vipp1_1′α6_L3_ and the equivalent in selected rungs of the Vipp1_HS_ C17 symmetry ring^1^. **D**, Hairpins slide relative to each other in Vipp1 ESCRT-III-like protofilaments. Two neighbouring subunits from rungs 1 and 4 of Vipp1_HS_ C14 ring are superposed using the right-hand subunits only (left and middle panels). Whilst Ala50 and Ala 141 are aligned in the superposed subunits (dark orange and blue), Ala141 is up to 14 Å apart between the neighbouring subunits (light orange and cyan; right panel) indicating hairpin sliding. The distances 41 Å and 61 Å (middle panel) relate to the typical span of minimum and maximum inter-ridge distances observed in Vipp1 rings_HS_, respectively. **E**, Lattice and subunit assignment of Vipp11′α6_1-219_ 2D planar filaments and sheets on a lipid monolayer, relating to Supplementary Figure 3I. The class average is shown with lattice points and dimensions based on the Fourier Transform (Supplementary Figure 3I) now assigned with Vipp1 ESCRT-III-like protofilaments positioned accordingly (orange and turquoise, equivalent to Bessel order n=17 in Vipp1_1′α6_L3_).

**Table S1.**
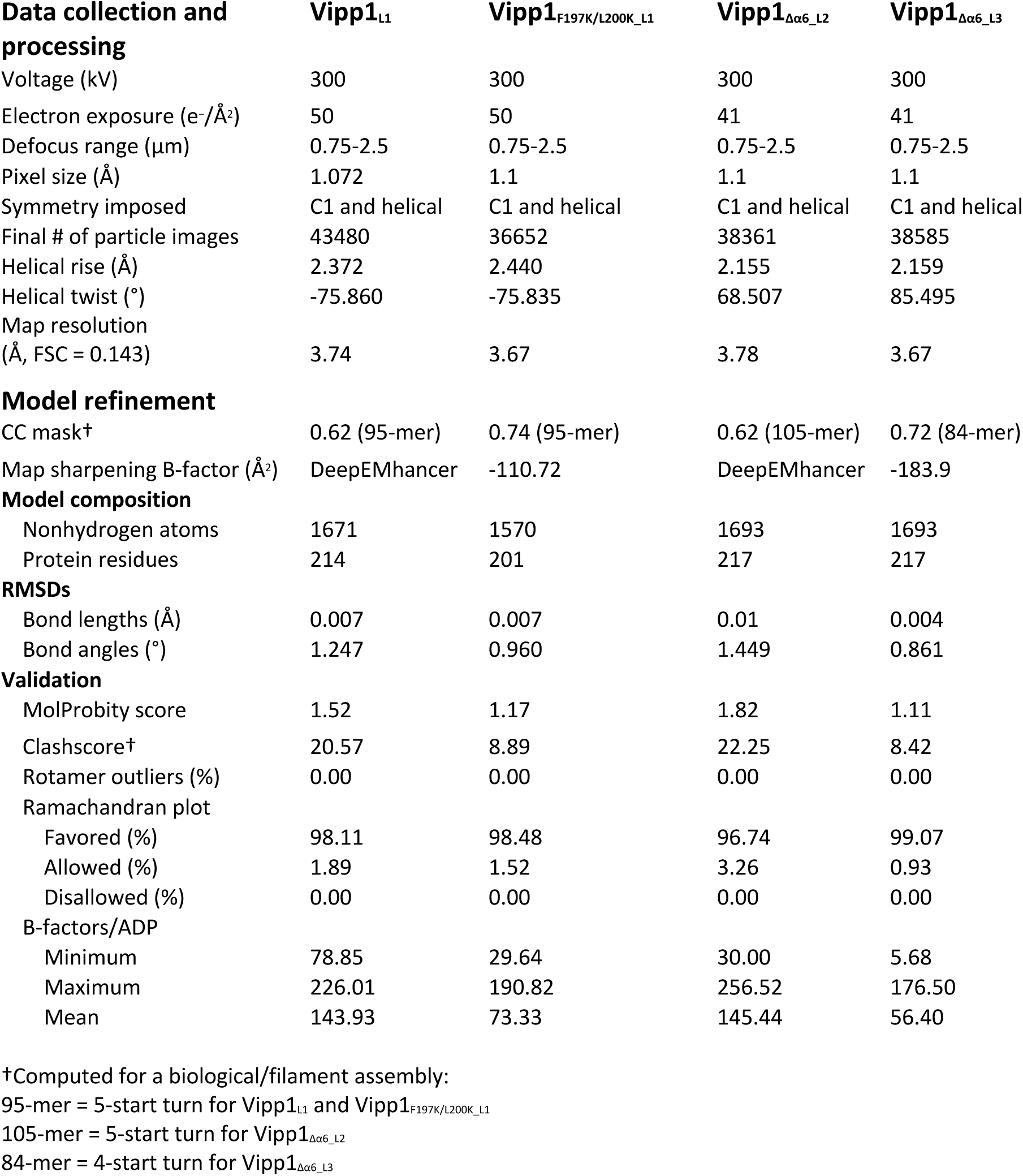
Cryo-EM data collection, refinement and validation statistics.

## Supplementary Videos

**Video S1 and S2, related to Figure 2C and Supplementary** Figure 1. F-AFM timecourse showing growth of Vipp1 planar filaments on a membrane patch.

**Video S3, related to Figure 3B.** F-AFM timecourse showing how spiral maturation and ring biogenesis correlated with increasing height offset between spiral inner turns and the membrane.

**Video S4, related to Supplementary** Figure 6F. Morph between constricted versus non-constricted ESCRT-III-like protofilaments in Vipp1_L1_ and Vipp1_1′α6_L3_. Five monomers were superposed with the alignment focussed on the central subunit.

**Video S5, related to Figure 6G.** Morph between superposed Vipp1_F197K/L200K_L1_ subunit with a Vipp1 ring_HS_ subunit (C14 symmetry, rung 4, PDB code 6ZW4).

## Methods

### Expression and purification of Vipp1 filament and ring assembly

All Vipp1 clones originated from a previous study^1^. In brief, the coding sequence for *N. punctiforme vipp1* (Uniprot code B2J6D9), *vipp1Δα6_1-219_* and *vipp1_F197K/L200K_* were cloned into pOPTM (a pET derivative) to yield an N-terminal MBP fusion with a TEV cleavage site in the linker. An N-terminal hexa-histidine tag was included on the MBP moiety. For the purification of all Vipp1 clones, plasmids were co-transformed into *E. coli* C43 (DE3) electro-competent cells (Lucigen) modified to incorporate a *pspA* gene knockout^60^. Cells were grown on LB-agarose with ampicillin (100 µg/ml). 2xYT media was inoculated and cells grown at 37°C until induction at OD600 = 0.6 with 1 mM isopropyl β-D-1-thiogalactopyranoside (IPTG). Cells were grown for 16 hours at 19°C and shaken at 200 rpm. All following steps were carried out at 4°C unless otherwise specified. Cell pellets were resuspended in buffer containing 50mM HEPES-NaOH pH 7.5, 500mM NaCl, 0.1mg/ml DNase-I and Roche cOmplete™ EDTA-free protease inhibitor cocktail and sonicated on ice. The lysate was clarified by centrifugation at 16,000 x g for 20 mins. The supernatant was incubated with 20 ml amylose resin (NEB) pre-equilibrated with lysis buffer for 15 mins before loading onto a gravity flow column. Resin was washed with three column volumes (CV) of wash buffer (50mM HEPES-NaOH pH 8.4, 200mM NaCl), two CV of ATP-wash buffer (50mM HEPES-NaOH pH 8.0, 80mM KCl, 2.5mM MgCl_2_ and 5mM ATP), followed by two CV of wash buffer. Sample was eluted with wash buffer supplemented with 15 mM maltose. Peak fractions were pooled and incubated with TEV initially at 34°C for two hours followed by 48 hours at room temperature. Digested samples were dialysed (12-14 kDa cut-off) overnight in size-exclusion chromatography (SEC) buffer containing 25mM HEPES-NaOH pH 8.4, 10 mM KCl, 10 mM MgCl_2_ except for Vipp1Δα6_1-219_ where 25mM HEPES-NaOH pH 8.4, 10mM NaCl was used. Samples were passed over a 10 ml Ni-NTA bead gravity flow column pre-equilibrated with SEC buffer to remove His-tagged MBP and TEV in the column. Vipp1 was collected from the flow through and concentrated using Vivaspin-20 concentrators with 10 kDa cut-off. During this process, sample was iteratively diluted with SEC buffer to avoid high protein concentrations in imidazole. Samples were injected onto a Sephacryl 16/60 S-500 pre-equilibrated with SEC buffer. Gel filtration yielded three peaks. The first peak after the void volume at ∼0.3 CV contained Vipp1 helical polymers used in this study, the second peak at ∼0.4-0.7 CV was predominantly populated with rings, and the third peak at ∼0.75-0.9 CV comprised Vipp1 monomer or small oligomers, MBP and TEV. For exclusive purification of Vipp1 rings_HS_, a protocol with high salt (50 mM) in the SEC buffer was used as described previously^1^.

### Lipid-covered silica beads preparation

For fluorescent light microscopy (FLM) studies, lipid lamellae were deposited on 40 µm silica beads as previously described^61^. Briefly, 1,2-dioleoyl-sn-glycero-3-phosphocholine (DOPC), 1,2-dioleoyl-sn-glycero-3-phospho-L-serine (DOPS), and (1,2-Dioleoyl-sn-glycero-3-phosphoethanolamine labeled with Atto 647N) Atto-647DOPE were mixed at a ratio of 59.95:40:0.05 mol%, respectively, from lipid stocks in chloroform to a final lipid concentration of 0.5 mg/ml. The lipid mixture was then dried in a vacuum for at least two hours to completely remove the chloroform, forming a dried lipid film, followed by hydration and resuspension in a buffer containing 1 mM HEPES-NaOH at pH 7.4. Subsequently, 1 µL of 40 µm silica beads (Microspheres-Nanospheres) was mixed with 10 µL of the hydrated lipid mixture and then divided into five drops placed on a parafilm surface. Subsequently, these beads-lipid drops were dried in a vacuum for at least 30 mins until the complete evaporation of the aqueous buffer.

### Supported lipid bilayers (SLBs) preparation

For FLM studies, SLBs were prepared as previously described^62^. Briefly, a coverslip was cleaned by sonication in water, ethanol, and water for 10 mins each, followed by 30 seconds of plasma cleaning (Harrick Plasma). After plasma cleaning, an Ibidi chamber (sticky-Slide VI 0.4) was mounted on the coverslip, and each of the wells was filled with 200 µL of buffer (25mM HEPES-NaOH, pH 8.3, 1mM EDTA, and 10 or 500 mM NaCl depending on the experiment). To form the SLBs, a portion of the lipid-covered silica beads was transferred to each well using a glass pipette, leading to the spilling of the lipid bilayers on the coverslip. The fluorescently labeled protein was introduced in the assays by replacing 100 µL buffer with new buffer containing the protein at desired concentration in the final volume of 200 µL.

### Large unilamellar vesicles (LUVs) preparatio

For F-AFM studies, LUVs were prepared using either *E. coli* total lipid extract (Avanti) or 1,2-dioleoyl-sn-glycero-3-phosphocholine (DOPC), 1,2-dioleoyl-sn-glycero-3-phospho-L-serine (DOPS) mixed at a ratio of 60:40 mol%. Both lipid mixes were tested with Vipp1 and similar polymers observed assembling including sheets, spirals and rings. Therefore, for the F-AFM results presented in this study *E. coli* total lipid extract was subsequently used to maintain consistency with EM monolayer studies. Lipids dissolved in a mixture of chloroform and methanol (1:1) were dried under N_2_ flux followed by overnight incubation in a vacuum oven at 37°C. Afterwards, lipids were fully rehydrated with buffer containing 25mM HEPES-NaOH pH 7.4 for 10 mins at room temperature, obtaining a 0.8 mg/mL lipid solution. The lipid suspension was vortexed for 30 seconds and freeze– thawed five times in liquid nitrogen and a water-bath. Mica-SLBs were then prepared by depositing LUVs onto freshly cleaved mica placed on the imaging chamber and incubating for 20 mins at 37°C with 10 mM HEPES-NaOH pH 7.4, 10mM CaCl_2_, and 2mM MgCl_2_. Samples were rinsed thoroughly by buffer containing 25mM HEPES-NaOH pH 7.4 with or without 150 mM NaCl for samples in high salt and low salt conditions, respectively. A final volume of 300µL of these buffers for their respective conditions was added into the imaging chamber.

### FLM image and movie acquisition

For the FLM studies, Vipp1 was chemically labelled at the N-terminus with TFP-AlexaFluor-488 (ThermoFisher Scientific) following the labelling procedure provided by the manufacturer. Fluorescence image acquisition was performed using an inverted spinning disc microscope assembled by 3i (Intelligent Imaging Innovation), consisting of a Nikon base (Eclipse C1, Nikon), a 100x 1.49 NA oil immersion objective, and an EVOLVE EM-CCD camera (Roper Scientific). The plugin Turboreg^63^ and a custom-written ImageJ macro were used for X-Y drift correction.

### Fast Scan AFM image acquisition

A JPK NanoWizard Ultraspeed AFM (Bruker and JPK BioAFM) equipped with USC-F0.3-k0.3-10 cantilevers with spring constant of 0.3N nm^−1^, resonance frequency of about 300 kHz (Nanoworld), was used for image acquisition. The F-AFM was operated in tapping mode, where the cantilever oscillated at a frequency proximal to 150kHz. Here, both topographic and phase images were reported. Initially, the supported lipid bilayer was imaged before selecting and imaging the area of interests (AOIs). While imaging of the AOIs was ongoing, Vipp1, Vipp1Δα6_1-219_ or Vipp1_F197K/L200K_ samples were injected into the imaging chamber at 3.5-7 µM or 21 µM for Vipp1 rings_HS_ in the final volume with either high or low salt buffer as described above in the LUV preparation. Images were analysed with JPKSPM Data Processing, ImageJ, and WSxM software^64^.

### Negative stain electron microscopy (NS EM) and data collection

To visualise Vipp1, 4 µL of 6 µM sample was spotted onto plasma cleaned carbon-coated EM grids (300-mesh, Agar Scientific) and incubated for 1 min. The samples were blotted, washed with water, and stained twice with two drops of 2 % uranyl acetate. Images were acquired on a FEI Tecnai 12 electron microscope equipped with a TVIPS 4K CMOS XF416 camera.

### Vipp1 monolayer assay and image analysis

Monolayer assays were performed following methods previously outlined^1^ but modified in buffer composition and lipid drop size. *E. coli* total lipid extract (Avanti Polar Lipids) was used to prepare the lipid monolayers. Wells (4 mm diameter) in a custom-built Teflon block were filled with 50 µL of assay buffer (25mM HEPES-NaOH, pH 8.4, 5mM KCl, and 5mM MgCl_2_) and a 3 µL drop of 0.1 mg/ml lipid dissolved in chloroform was gently applied on top and allowed to evaporate for one hour. Carbon-coated EM grids (200-mesh, Agar Scientific) without plasma cleaning or glow discharge were placed on the lipid layer with the carbon side facing the lipid. 15 µM Vipp1 was added through a side port and mixed gently with the buffer beneath the lipid layer. Control wells without protein or with protein-only (chloroform drop without lipid) were included. The assays were incubated for one hour, and grids were subsequently removed, stained and imaged on a Philips Tecnai 12 electron microscope. Datasets containing 136 and 139 images for Vipp1 and Vipp1Δα6_1-219_, respectively, were collected at 2.563 Å pixel size. The contrast transfer function (CTF) was estimated with CTFFIND-4.1^65^. Filament segments were picked and extracted for three rounds of 2D classification in Relion4^66^. The final 2D class averages presented incorporated 1951 and 6902 segments for Vipp1 and Vipp1Δα6_1-219_, respectively.

### Cryo-EM sample preparation and data collection

3.5 µL of Vipp1 at a final concentration of 60 µM was incubated for 90 seconds on a plasma cleaned holey R2/2 Quantifoil copper grid (Electron Microscopy Science) before plunge freezing in liquid ethane using a Vitrobot Mark IV (FEI) set at 100 % humidity and 10°C. Cryo-EM data was collected at 300 kV on a Titan Krios equipped with a Gatan Quantum K2 Summit detector operated in super resolution mode with a pixel size of 0.536 Å. A total of 19,740 movies were acquired at a defocus range between 0.75 and 2.5 µM with a total electron dose of 50 e^-^/Å^2^ fractionated over 50 frames and an exposure of 4.1 seconds. For Vipp1_F197K/L200K_ and Vipp1Δα6_1-219_, cryo-EM data was collected at 300 kV on a Titan Krios microscope (LonCEM, The Francis Crick Institute, London, UK) equipped with a Gatan K3 detector operated in super resolution mode with a pixel size of 0.55 Å. A total of 30,216 and 29,562 micrographs were acquired at a defocus range between 0.75 and 2.5 µM with a total electron dose of 50 and 41 e^-^/Å^2^ fractionated over 44 and 28 frames using 4.6 and 3 second exposures, respectively.

### Cryo-EM image processing and helical reconstruction of Vipp1 filaments

Micrograph movies were corrected for beam-induced sample motion and Fourier cropped to a pixel size of 1.072 Å for Vipp1, or 1.1 Å for Vipp1_F197K/L200K_ and Vipp1Δα6_1-219_ using MotionCor2^67^. Contrast-transfer function (CTF) estimations were performed using CTFFIND-4.1. Filaments were boxed into overlapping particles using crYOLO^68^. Each particle box overlapped with its neighbour by 46 Å (Vipp1_L1_, Vipp1_F197K/L200K_L1_, Vipp1_Δα6_L2_) or 44 Å (Vipp1_Δα6_L3_) equivalent to the left-handed 5-start or 4-start helical pitches, respectively. Using Relion4, this resulted in extraction of 138562, 1172337 and 648059 particles with 504 pixel box size for Vipp1, Vipp1_F197K/L200K_ and Vipp1Δα6_1-219_, respectively. Particles were then binned by a factor of three and imported into cryoSPARC^69^ for iterative rounds of 2D classification. Low quality particles were discarded, and remaining particles sorted into separate bins based on their Fourier Transform. Attempts to determine helical symmetry parameters based on C1 reconstructions yielded only low quality reconstructions without meaningful symmetries. Only Vipp1_Δα6_L3_ symmetry particle class averages gave a Fourier Transform with non-overlapping layer lines amenable to indexing (Supplementary Figure 4). This gave a grid of possible symmetries which were systematically tested using helical refinement in Cryosparc. The helical parameters that yielded a reconstruction showing obvious secondary structure features were used for next steps in Relion4. This reconstruction also served as a subsequent initial reference map. For all other symmetries including Vipp1_L1_, Vipp1_F197K/L200K_L1_ and Vipp1_Δα6_L2_, particle class averages gave Fourier Transforms with overlapping layer lines that impeded indexing. Initial helical parameters were therefore determined by calculating and screening a grid of theoretical lattices close to Vipp1_1′α6_L3_ symmetry. Cleaned stacks containing 64936, 508377, 91126 and 57329 particles relating to Vipp1_L1_, Vipp1_F197K/L200K_L1_, Vipp1_Δα6_L2_ and Vipp1_Δα6_L3_ were exported from Cryosparc and re-extracted in Relion using a 150 pixel box size so that particles remained binned by a factor of three. Iterative rounds of 3D classification were undertaken resulting in final particle stacks of 43480, 36652, 38361 and 38585 relating to Vipp1_L1_, Vipp1_F197K/L200K_L1_, Vipp1_Δα6_L2_ and Vipp1_Δα6_L3_. Particles were re-extracted with no binning using a 450 pixel box size for iterative rounds of 3D autorefinement incorporating three rounds of CTF refinement and Bayesian polishing. For these steps a mask covering the central 30 % of the map was used. Final refinements for Vipp1_L1_, Vipp1_F197K/L200K_L1_, Vipp1_Δα6_L2_ and Vipp1_Δα6_L3_ converged with helical rises of 2.372, 2.440, 2.155 and 2.159 Å, and helical twists of -75.860°, -75.835°, 68.507° and 85.495°, respectively. Final resolutions of 3.7, 3.7, 3.8 and 3.7 Å were based on gold standard Fourier shell correlations (FSC) = 0.143. Vipp1_L1_ and Vipp1_Δα6_L2_ maps were sharpened using DeepEMhancer^59^ wideTarget model, Vipp1_F197K/L200K_L1_ was sharpened using Relion4 postprocess with B-factor -110.719 Å^2^, whilst Vipp1_Δα6_L3_ used Phenix^70^ auto_sharpen with B-factor -183.9 Å^2^. Map local resolution was estimated using ResMap^71^.

### Model building and refinement

A monomer extracted from Vipp1 C14 symmetry ring (PDB code 6ZW4) was fitted into Vipp1_Δα6_L3_ as an initial build template. High quality map with generally excellent side chain detail facilitated manual building and modelling in Coot^72^ and ISOLDE^73^. By applying the helical parameters to this asymmetric unit, a helical filament was generated in ChimeraX 1.4^74^. A central subunit was chosen and all subunits that were not within 5 Å of this central subunit deleted. This assembly that comprised 19 subunits (termed 19-mer) was then used for model building with the central subunit now modelled in the context of its neighbours. At the end of this iteration, the central subunit was extracted and a new helical filament and 19-mer generated. This 19-mer was used for real-space refinement in Phenix after which a new 19-mer was generated again using the central subunit. This model building process was iterated with the same workflow utilised for Vipp1_L1_, Vipp1_F197K/L200K_L1_ and Vipp1_Δα6_L2_. In the lower resolution regions of Vipp1_L1_ and Vipp1_Δα6_L2_ maps where side chain detail was reduced or absent, side chains were modelled based on Vipp1_Δα6_L3_ and Vipp1_F197K/L200K_L1_ structures. Final refinement and model validation statistics are in Table S1.

### Cryo-ET sample preparation, data collection and image processing

5 nm gold fiducials without BSA (Sigma-Aldrich) were mixed with 60 µM Vipp1Δα6_1-219_ sample. 3.5 µL of this mix was incubated on plasma cleaned holey R2/2 Quantifoil copper 200 mesh grids for 90 seconds at 10°C and vitrified as for cryo-EM samples. Tomograms were collected at 300 kV on a Titan Krios electron microscope (LonCEM, UK) equipped with a Gatan K3 detector with pixel size of 2.257 Å. Dose-symmetric tilt series were acquired between -60° to +60° with 3° intervals and a defocus range between 3-6 µm. A total accumulated dose of 101.6 e^-^/Å^2^ was fractionated over 15 frames per tilt using an exposure of 5.6 seconds. Tilt series movies were corrected for beam-induced sample motion using MotionCor2^67^. Using IMOD^75^, the tilt-series was 5-binned to yield a pixel size = 11.3 Å, aligned and reconstructed using a SIRT algorithm.

## Data and code availability

3D cryo-EM density maps produced in this study have been deposited in the Electron Microscopy Data Bank with accession code EMD-18318, EMD-18319, EMD-18321 and EMD-18322 for Vipp1_L1_, Vipp1_F197K/L200K_L1_, Vipp1_1′α6_L2_ and Vipp1_1′α6_L3_, respectively. Atomic coordinates have been deposited in the Protein Data Bank (PDB) under accession code 8QBR, 8QBS, 8QBV and 8QBW, respectively.

## References

1. Liu, J., Tassinari, M., Souza, D.P., Naskar, S., Noel, J.K., Bohuszewicz, O., Buck, M., Williams, T.A., Baum, B., and Low, H.H. (2021). Bacterial Vipp1 and PspA are members of the ancient ESCRT-III membrane-remodeling superfamily. Cell 184, 3660–3673.e18. 10.1016/j.cell.2021.05.041.

2. Tarrason Risa, G., Hurtig, F., Bray, S., Hafner, A.E., Harker-Kirschneck, L., Faull, P., Davis, C., Papatziamou, D., Mutavchiev, D.R., Fan, C., et al. (2020). The proteasome controls ESCRT-III–mediated cell division in an archaeon. Science (1979) 369. 10.1126/science.aaz2532.

3. Votteler, J., and Sundquist, W.I. (2013). Virus Budding and the ESCRT Pathway. Cell Host Microbe 14, 232–241. 10.1016/j.chom.2013.08.012.

4. Liu, J., Gao, R., Li, C., Ni, J., Yang, Z., Zhang, Q., Chen, H., and Shen, Y. (2017). Functional assignment of multiple ESCRT-III homologs in cell division and budding in *Sulfolobus islandicus*. Mol Microbiol 105, 540–553. 10.1111/mmi.13716.

5. Juan, T., and Fürthauer, M. (2018). Biogenesis and function of ESCRT-dependent extracellular vesicles. Semin Cell Dev Biol 74, 66–77. 10.1016/j.semcdb.2017.08.022.

6. Ellen, A.F., Albers, S.-V., Huibers, W., Pitcher, A., Hobel, C.F. V., Schwarz, H., Folea, M., Schouten, S., Boekema, E.J., Poolman, B., et al. (2009). Proteomic analysis of secreted membrane vesicles of archaeal Sulfolobus species reveals the presence of endosome sorting complex components. Extremophiles 13, 67–79. 10.1007/s00792-008-0199-x.

7. Tang, S., Henne, W.M., Borbat, P.P., Buchkovich, N.J., Freed, J.H., Mao, Y., Fromme, J.C., and Emr, S.D. (2015). Structural basis for activation, assembly and membrane binding of ESCRT-III Snf7 filaments. Elife 4. 10.7554/eLife.12548.

8. Olmos, Y. (2022). The ESCRT Machinery: Remodeling, Repairing, and Sealing Membranes. Membranes (Basel) 12, 633. 10.3390/membranes12060633.

9. Junglas, B., Huber, S.T., Heidler, T., Schlösser, L., Mann, D., Hennig, R., Clarke, M., Hellmann, N., Schneider, D., and Sachse, C. (2021). PspA adopts an ESCRT-III-like fold and remodels bacterial membranes. Cell 184, 3674–3688.e18. 10.1016/j.cell.2021.05.042.

10. Brissette, J.L., Russel, M., Weiner, L., and Model, P. (1990). Phage shock protein, a stress protein of Escherichia coli. Proc Natl Acad Sci U S A 87. 10.1073/pnas.87.3.862.

11. Joly, N., Engl, C., Jovanovic, G., Huvet, M., Toni, T., Sheng, X., Stumpf, M.P.H., and Buck, M. (2010). Managing membrane stress: the phage shock protein (Psp) response, from molecular mechanisms to physiology. FEMS Microbiol Rev 34, 797–827. 10.1111/j.1574-6976.2010.00240.x.

12. Kobayashi, R., Suzuki, T., and Yoshida, M. (2007). Escherichia coli phage-shock protein A (PspA) binds to membrane phospholipids and repairs proton leakage of the damaged membranes. Mol Microbiol 66, 100–109. 10.1111/j.1365-2958.2007.05893.x.

13. McDonald, C., Jovanovic, G., Ces, O., and Buck, M. (2015). Membrane Stored Curvature Elastic Stress Modulates Recruitment of Maintenance Proteins PspA and Vipp1. mBio 6. 10.1128/mBio.01188-15.

14. Yamaguchi, S., Gueguen, E., Horstman, N.K., and Darwin, A.J. (2010). Membrane association of PspA depends on activation of the phage-shock-protein response in Yersinia enterocolitica. Mol Microbiol 78, 429–443. 10.1111/j.1365-2958.2010.07344.x.

15. Aseeva, E., Ossenbühl, F., Sippel, C., Cho, W.K., Stein, B., Eichacker, L.A., Meurer, J., Wanner, G., Westhoff, P., Soll, J., et al. (2007). Vipp1 is required for basic thylakoid membrane formation but not for the assembly of thylakoid protein complexes. Plant Physiology and Biochemistry 45, 119–128. 10.1016/j.plaphy.2007.01.005.

16. Fuhrmann, E., Gathmann, S., Rupprecht, E., Golecki, J., and Schneider, D. (2009). Thylakoid Membrane Reduction Affects the Photosystem Stoichiometry in the Cyanobacterium *Synechocystis* sp. PCC 6803. Plant Physiol 149, 735–744. 10.1104/pp.108.132373.

17. Gao, H., and Xu, X. (2009). Depletion of Vipp1 in *Synechocystis* sp. PCC 6803 affects photosynthetic activity before the loss of thylakoid membranes. FEMS Microbiol Lett 292, 63–70. 10.1111/j.1574-6968.2008.01470.x.

18. Gutu, A., Chang, F., and O’Shea, E.K. (2018). Dynamical localization of a thylakoid membrane binding protein is required for acquisition of photosynthetic competency. Mol Microbiol 108, 16–31. 10.1111/mmi.13912.

19. Nordhues, A., Schöttler, M.A., Unger, A.-K., Geimer, S., Schönfelder, S., Schmollinger, S., Rütgers, M., Finazzi, G., Soppa, B., Sommer, F., et al. (2012). Evidence for a Role of VIPP1 in the Structural Organization of the Photosynthetic Apparatus in *Chlamydomonas*. Plant Cell 24, 637–659. 10.1105/tpc.111.092692.

20. Kroll, D., Meierhoff, K., Bechtold, N., Kinoshita, M., Westphal, S., Vothknecht, U.C., Soll, J., and Westhoff, P. (2001). *VIPP1*, a nuclear gene of *Arabidopsis thaliana* essential for thylakoid membrane formation. Proceedings of the National Academy of Sciences 98, 4238–4242. 10.1073/pnas.061500998.

21. Lo, S.M., and Theg, S.M. (2012). Role of Vesicle-Inducing Protein in Plastids 1 in cpTat transport at the thylakoid. The Plant Journal 71, 656–668. 10.1111/j.1365-313X.2012.05020.x.

22. Walter, B., Hristou, A., Nowaczyk, M.M., and Schünemann, D. (2015). *In vitro* reconstitution of co-translational D1 insertion reveals a role of the cpSec–Alb3 translocase and Vipp1 in photosystem II biogenesis. Biochemical Journal 468, 315–324. 10.1042/BJ20141425.

23. Westphal, S., Heins, L., Soll, J., and Vothknecht, U.C. (2001). *Vipp1* deletion mutant of *Synechocystis* : A connection between bacterial phage shock and thylakoid biogenesis? Proceedings of the National Academy of Sciences 98, 4243–4248. 10.1073/pnas.061501198.

24. Zhang, L., and Sakamoto, W. (2015). Possible function of VIPP1 in maintaining chloroplast membranes. Biochimica et Biophysica Acta (BBA) - Bioenergetics 1847, 831–837. 10.1016/j.bbabio.2015.02.013.

25. McCullough, J., Frost, A., and Sundquist, W.I. (2018). Structures, Functions, and Dynamics of ESCRT-III/Vps4 Membrane Remodeling and Fission Complexes. Annu Rev Cell Dev Biol 34, 85–109. 10.1146/annurev-cellbio-100616-060600.

26. Pfitzner, A.-K., Moser von Filseck, J., and Roux, A. (2021). Principles of membrane remodeling by dynamic ESCRT-III polymers. Trends Cell Biol 31, 856–868. 10.1016/j.tcb.2021.04.005.

27. Buchkovich, N.J., Henne, W.M., Tang, S., and Emr, S.D. (2013). Essential N-Terminal insertion motif anchors the ESCRT-III filament during MVB vesicle formation. Dev Cell 27. 10.1016/j.devcel.2013.09.009.

28. Otters, S., Braun, P., Hubner, J., Wanner, G., Vothknecht, U.C., and Chigri, F. (2013). The first α-helical domain of the vesicle-inducing protein in plastids 1 promotes oligomerization and lipid binding. Planta 237, 529–540. 10.1007/s00425-012-1772-1.

29. Heidrich, J., Wulf, V., Hennig, R., Saur, M., Markl, J., Sönnichsen, C., and Schneider, D. (2016). Organization into higher ordered ring structures counteracts membrane binding of IM30, a protein associated with inner membranes in chloroplasts and cyanobacteria. Journal of Biological Chemistry 291. 10.1074/jbc.M116.722686.

30. Zhang, L., Kondo, H., Kamikubo, H., Kataoka, M., and Sakamoto, W. (2016). VIPP1 Has a Disordered C-Terminal Tail Necessary for Protecting Photosynthetic Membranes against Stress. Plant Physiol 171, 1983–1995. 10.1104/pp.16.00532.

31. Hennig, R., West, A., Debus, M., Saur, M., Markl, J., Sachs, J.N., and Schneider, D. (2017). The IM30/Vipp1 C-terminus associates with the lipid bilayer and modulates membrane fusion. Biochimica et Biophysica Acta (BBA) - Bioenergetics 1858, 126–136. 10.1016/j.bbabio.2016.11.004.

32. Hennig, R., Heidrich, J., Saur, M., Schmüser, L., Roeters, S.J., Hellmann, N., Woutersen, S., Bonn, M., Weidner, T., Markl, J., et al. (2015). IM30 triggers membrane fusion in cyanobacteria and chloroplasts. Nat Commun 6, 7018. 10.1038/ncomms8018.

33. McCullough, J., Clippinger, A.K., Talledge, N., Skowyra, M.L., Saunders, M.G., Naismith, T. V., Colf, L.A., Afonine, P., Arthur, C., Sundquist, W.I., et al. (2015). Structure and membrane remodeling activity of ESCRT-III helical polymers. Science (1979) 350, 1548–1551. 10.1126/science.aad8305.

34. Gupta, T.K., Klumpe, S., Gries, K., Heinz, S., Wietrzynski, W., Ohnishi, N., Niemeyer, J., Spaniol, B., Schaffer, M., Rast, A., et al. (2021). Structural basis for VIPP1 oligomerization and maintenance of thylakoid membrane integrity. Cell 184, 3643–3659.e23. 10.1016/j.cell.2021.05.011.

35. Cashikar, A.G., Shim, S., Roth, R., Maldazys, M.R., Heuser, J.E., and Hanson, P.I. (2014). Structure of cellular ESCRT-III spirals and their relationship to HIV budding. Elife 3. 10.7554/eLife.02184.

36. Effantin, G., Dordor, A., Sandrin, V., Martinelli, N., Sundquist, W.I., Schoehn, G., and Weissenhorn, W. (2013). ESCRT-III CHMP2A and CHMP3 form variable helical polymers in vitro and act synergistically during HIV-1 budding. Cell Microbiol 15. 10.1111/cmi.12041.

37. Pires, R., Hartlieb, B., Signor, L., Schoehn, G., Lata, S., Roessle, M., Moriscot, C., Popov, S., Hinz, A., Jamin, M., et al. (2009). A Crescent-Shaped ALIX Dimer Targets ESCRT-III CHMP4 Filaments. Structure 17. 10.1016/j.str.2009.04.007.

38. Dobro, M.J., Samson, R.Y., Yu, Z., McCullough, J., Ding, H.J., Chong, P.L.-G., Bell, S.D., and Jensen, G.J. (2013). Electron cryotomography of ESCRT assemblies and dividing *Sulfolobus* cells suggests that spiraling filaments are involved in membrane scission. Mol Biol Cell 24, 2319–2327. 10.1091/mbc.e12-11-0785.

39. Henne, W.M., Buchkovich, N.J., Zhao, Y., and Emr, S.D. (2012). The Endosomal Sorting Complex ESCRT-II Mediates the Assembly and Architecture of ESCRT-III Helices. Cell 151, 356–371. 10.1016/j.cell.2012.08.039.

40. Chiaruttini, N., Redondo-Morata, L., Colom, A., Humbert, F., Lenz, M., Scheuring, S., and Roux, A. (2015). Relaxation of Loaded ESCRT-III Spiral Springs Drives Membrane Deformation. Cell 163, 866–879. 10.1016/j.cell.2015.10.017.

41. Mierzwa, B.E., Chiaruttini, N., Redondo-Morata, L., Moser von Filseck, J., König, J., Larios, J., Poser, I., Müller-Reichert, T., Scheuring, S., Roux, A., et al. (2017). Dynamic subunit turnover in ESCRT-III assemblies is regulated by Vps4 to mediate membrane remodelling during cytokinesis. Nat Cell Biol 19, 787–798. 10.1038/ncb3559.

42. Shen, Q.-T., Schuh, A.L., Zheng, Y., Quinney, K., Wang, L., Hanna, M., Mitchell, J.C., Otegui, M.S., Ahlquist, P., Cui, Q., et al. (2014). Structural analysis and modeling reveals new mechanisms governing ESCRT-III spiral filament assembly. Journal of Cell Biology 206, 763–777. 10.1083/jcb.201403108.

43. Azad, K., Guilligay, D., Boscheron, C., Maity, S., De Franceschi, N., Sulbaran, G., Effantin, G., Wang, H., Kleman, J.-P., Bassereau, P., et al. (2023). Structural basis of CHMP2A– CHMP3 ESCRT-III polymer assembly and membrane cleavage. Nat Struct Mol Biol 30, 81–90. 10.1038/s41594-022-00867-8.

44. Hanson, P.I., Roth, R., Lin, Y., and Heuser, J.E. (2008). Plasma membrane deformation by circular arrays of ESCRT-III protein filaments. Journal of Cell Biology 180. 10.1083/jcb.200707031.

45. Nguyen, H.C., Talledge, N., McCullough, J., Sharma, A., Moss, F.R., Iwasa, J.H., Vershinin, M.D., Sundquist, W.I., and Frost, A. (2020). Membrane constriction and thinning by sequential ESCRT-III polymerization. Nat Struct Mol Biol 27, 392–399. 10.1038/s41594-020-0404-x.

46. Junglas, B., Orru, R., Axt, A., Siebenaller, C., Steinchen, W., Heidrich, J., Hellmich, U.A., Hellmann, N., Wolf, E., Weber, S.A.L., et al. (2020). IM30 IDPs form a membrane-protective carpet upon super-complex disassembly. Commun Biol 3. 10.1038/s42003-020-01314-4.

47. Chiaruttini, N., and Roux, A. (2017). Dynamic and elastic shape transitions in curved ESCRT-III filaments. Curr Opin Cell Biol 47, 126–135. 10.1016/j.ceb.2017.07.002.

48. Harker-Kirschneck, L., Baum, B., and Šarić, A. ela (2019). Changes in ESCRT-III filament geometry drive membrane remodelling and fission in silico. BMC Biol 17. 10.1186/s12915-019-0700-2.

49. Pfitzner, A.-K., Mercier, V., Jiang, X., Moser von Filseck, J., Baum, B., Šarić, A., and Roux, A. (2020). An ESCRT-III Polymerization Sequence Drives Membrane Deformation and Fission. Cell 182, 1140–1155.e18. 10.1016/j.cell.2020.07.021.

50. Moser von Filseck, J., Barberi, L., Talledge, N., Johnson, I.E., Frost, A., Lenz, M., and Roux, A. (2020). Anisotropic ESCRT-III architecture governs helical membrane tube formation. Nat Commun 11, 1516. 10.1038/s41467-020-15327-4.

51. Jukic, N., Perrino, A.P., Humbert, F., Roux, A., and Scheuring, S. (2022). Snf7 spirals sense and alter membrane curvature. Nat Commun 13, 2174. 10.1038/s41467-022-29850-z.

52. Diaz, R., Rice, William, J., and Stokes, D.L. (2010). Fourier–Bessel Reconstruction of Helical Assemblies. In Bone, pp. 131–165. 10.1016/S0076-6879(10)82005-1.

53. Theis, J., Gupta, T.K., Klingler, J., Wan, W., Albert, S., Keller, S., Engel, B.D., and Schroda, M. (2019). VIPP1 rods engulf membranes containing phosphatidylinositol phosphates. Sci Rep 9, 8725. 10.1038/s41598-019-44259-3.

54. Sun, C., Gonzalez, B., and Jiang, W. (2022). Helical Indexing in Real Space. Sci Rep 12, 8162. 10.1038/s41598-022-11382-7.

55. Ohnishi, N., Zhang, L., and Sakamoto, W. (2018). VIPP1 Involved in Chloroplast Membrane Integrity Has GTPase Activity in Vitro. Plant Physiol 177, 328–338. 10.1104/pp.18.00145.

56. Zhang, S., Shen, G., Li, Z., Golbeck, J.H., and Bryant, D.A. (2014). Vipp1 Is Essential for the Biogenesis of Photosystem I but Not Thylakoid Membranes in Synechococcus sp. PCC 7002. Journal of Biological Chemistry 289, 15904–15914. 10.1074/jbc.M114.555631.

57. Rütgers, M., and Schroda, M. (2013). A role of VIPP1 as a dynamic structure within thylakoid centers as sites of photosystem biogenesis? Plant Signal Behav 8, e27037. 10.4161/psb.27037.

58. Kleerebezem, M., Crielaard, W., and Tommassen, J. (1996). Involvement of stress protein PspA (phage shock protein A) of Escherichia coli in maintenance of the protonmotive force under stress conditions. EMBO J 15, 162–171. 10.1002/j.1460-2075.1996.tb00344.x.

59. Sanchez-Garcia, R., Gomez-Blanco, J., Cuervo, A., Carazo, J.M., Sorzano, C.O.S., and Vargas, J. (2021). DeepEMhancer: a deep learning solution for cryo-EM volume post-processing. Commun Biol 4, 874. 10.1038/s42003-021-02399-1.

60. Chernyatina, A.A., and Low, H.H. (2019). Core architecture of a bacterial type II secretion system. Nat Commun 10, 5437. 10.1038/s41467-019-13301-3.

61. Velasco-Olmo, A., Ormaetxea Gisasola, J., Martinez Galvez, J.M., Vera Lillo, J., and Shnyrova, A. V. (2019). Combining patch-clamping and fluorescence microscopy for quantitative reconstitution of cellular membrane processes with Giant Suspended Bilayers. Sci Rep 9, 7255. 10.1038/s41598-019-43561-4.

62. Pucadyil, T.J., and Schmid, S.L. (2010). Supported Bilayers with Excess Membrane Reservoir: A Template for Reconstituting Membrane Budding and Fission. Biophys J 99, 517–525. 10.1016/j.bpj.2010.04.036.

63. Thevenaz, P., Ruttimann, U.E., and Unser, M. (1998). A pyramid approach to subpixel registration based on intensity. IEEE Transactions on Image Processing 7, 27–41. 10.1109/83.650848.

64. Horcas, I., Fernández, R., Gómez-Rodríguez, J.M., Colchero, J., Gómez-Herrero, J., and Baro, A.M. (2007). WSXM : A software for scanning probe microscopy and a tool for nanotechnology. Review of Scientific Instruments 78. 10.1063/1.2432410.

65. Rohou, A., and Grigorieff, N. (2015). CTFFIND4: Fast and accurate defocus estimation from electron micrographs. J Struct Biol 192, 216–221. 10.1016/j.jsb.2015.08.008.

66. Kimanius, D., Dong, L., Sharov, G., Nakane, T., and Scheres, S.H.W. (2021). New tools for automated cryo-EM single-particle analysis in RELION-4.0. Biochemical Journal 478, 4169–4185. 10.1042/BCJ20210708.

67. Zheng, S.Q., Palovcak, E., Armache, J.-P., Verba, K.A., Cheng, Y., and Agard, D.A. (2017). MotionCor2: anisotropic correction of beam-induced motion for improved cryo-electron microscopy. Nat Methods 14, 331–332. 10.1038/nmeth.4193.

68. Wagner, T., Merino, F., Stabrin, M., Moriya, T., Antoni, C., Apelbaum, A., Hagel, P., Sitsel, O., Raisch, T., Prumbaum, D., et al. (2019). SPHIRE-crYOLO is a fast and accurate fully automated particle picker for cryo-EM. Commun Biol 2, 218. 10.1038/s42003-019-0437-z.

69. Punjani, A., Rubinstein, J.L., Fleet, D.J., and Brubaker, M.A. (2017). cryoSPARC: algorithms for rapid unsupervised cryo-EM structure determination. Nat Methods 14, 290–296. 10.1038/nmeth.4169.

70. Liebschner, D., Afonine, P. V., Baker, M.L., Bunkóczi, G., Chen, V.B., Croll, T.I., Hintze, B., Hung, L.-W., Jain, S., McCoy, A.J., et al. (2019). Macromolecular structure determination using X-rays, neutrons and electrons: recent developments in *Phenix*. Acta Crystallogr D Struct Biol 75, 861–877. 10.1107/S2059798319011471.

71. Kucukelbir, A., Sigworth, F.J., and Tagare, H.D. (2014). Quantifying the local resolution of cryo-EM density maps. Nat Methods 11, 63–65. 10.1038/nmeth.2727.

72. Emsley, P., Lohkamp, B., Scott, W.G., and Cowtan, K. (2010). Features and development of *Coot*. Acta Crystallogr D Biol Crystallogr 66, 486–501. 10.1107/S0907444910007493.

73. Croll, T.I. (2018). *ISOLDE* : a physically realistic environment for model building into low-resolution electron-density maps. Acta Crystallogr D Struct Biol 74, 519–530. 10.1107/S2059798318002425.

74. Pettersen, E.F., Goddard, T.D., Huang, C.C., Meng, E.C., Couch, G.S., Croll, T.I., Morris, J.H., and Ferrin, T.E. (2021). UCSF ChimeraX : Structure visualization for researchers, educators, and developers. Protein Science 30, 70–82. 10.1002/pro.3943.

75. Kremer, J.R., Mastronarde, D.N., and McIntosh, J.R. (1996). Computer Visualization of Three-Dimensional Image Data Using IMOD. J Struct Biol 116, 71–76. 10.1006/jsbi.1996.0013.

